# Restricting Dietary Isoleucine Promotes Foxo3-Dependent Mitochondrial Respiration by Reducing Caloric Intake

**DOI:** 10.64898/2026.07.01.735791

**Authors:** Julianne Austin, Minzhen He, Zhi Yang, Dayyan Sayed, Danish Sayed, Maha Abdellatif

## Abstract

Our prior work demonstrated that lowering dietary branched-chain amino acids (BCAAs) improves cardiac outcomes during pressure overload-induced stress. Here, we identify isoleucine restriction (IleR) as the key driver of this effect. Dietary isoleucine restriction induces hypophagia and weight loss, recapitulating the effects of caloric restriction (CR). Although it does not prevent the initial development of left ventricular hypertrophy, it halts its progression and the decline in ejection fraction compared with controls. This is associated with preservation of electron transport chain (ETC) gene expression, cristae structure, NAD^+^/NADH levels, and mitochondrial respiratory capacity in cardiomyocytes, which is recapitulated by CR. Mechanistically, both IleR and CR diets increase Foxo3 expression, thereby blocking the decline in expression of its target ETC and mitochondrial genome-encoded genes. Consequently, this improves mitochondrial respiratory capacity and reduces cardiac fibrosis. We conclude that restricting dietary isoleucine improves cardiac health by increasing Foxo3 expression and mitochondrial function via a cell-autonomous mechanism and by reducing caloric intake.

## Introduction

Metabolic stress refers to the cell’s inability to meet its energy demands. This could be triggered by insufficient (e.g., starvation) or excessive (e.g., hyperglycemia or hyperlipidemia) nutrients. Also, increased mechanical stress (e.g., excessive or prolonged work overload) and ischemia trigger metabolic stress in the heart, as energy demand exceeds supply (Li et al., 2025). Initially, the heart can adapt by increasing energy output to meet demand, but if the demand is excessive or prolonged, as in disease states, energy supply is compromised by a decline in NAD^+^, FAD^+^, mitochondrial enzymes, and ETC subunits, as observed in heart failure (Lopaschuk et al., 2021). We have reported that we can reduce metabolic stress during increased pressure overload on the heart by restricting dietary branched-chain amino acids (isoleucine, leucine, and valine) in mice, thereby decelerating the transition from cardiac hypertrophy to heart failure (Yang et al., 2023).

Other dietary interventions that are powerful modulators of metabolic health and cardiovascular function include calorie restriction (CR). CR has been extensively studied for its ability to extend lifespan and improve cardiometabolic outcomes across species (Lin et al., 2002; Nicolas et al., 1999). The beneficial effects of CR are mediated, in part, by transcriptional and metabolic adaptations that enhance mitochondrial function, reduce oxidative stress, and improve cellular resilience (Gonzalez et al., 2004; Linford et al., 2007; Nisoli et al., 2005). A key mediator of these effects is the FOXO family of transcription factors, especially FOXO3a, which regulates genes involved in oxidative metabolism, stress resistance, and mitochondrial homeostasis (Kim et al., 2014; Makino et al., 2016; Qin et al., 2008; Wang et al., 2007). Activation of FOXO3a during nutrient limitation promotes the expression of oxidative phosphorylation genes and supports mitochondrial integrity, thereby preserving cellular energetics under stress conditions (Chaanine et al., 2016; Peserico et al., 2013).

Beyond overall caloric intake, emerging evidence suggests that the composition of dietary macronutrients—particularly amino acids—plays a crucial role in determining metabolic outcomes. Protein restriction and selective amino acid restriction have both been shown to improve metabolic health and extend lifespan in animal models. Notably, recent studies demonstrate that restricting the single BCAA, isoleucine, is sufficient to recapitulate many of the benefits of low-protein or low-BCAA diets, including improved metabolic health, reduced adiposity, and enhanced longevity (Green et al., 2023). These findings highlight isoleucine as a key dietary determinant of systemic metabolism and suggest that individual amino acids may exert distinct and non-redundant effects on physiological processes.

Despite these advances, the role of specific BCAAs, particularly isoleucine, in regulating cardiac remodeling and mitochondrial function under conditions of hemodynamic stress remains incompletely understood. Pressure overload, a common experimental and clinical model of cardiac hypertrophy and heart failure, is associated with mitochondrial dysfunction, impaired oxidative phosphorylation, and alterations in NAD⁺ redox balance, all of which contribute to disease progression (Lee et al., 2016). Furthermore, cardiac fibrosis—a key determinant of adverse remodeling—is closely linked to mitochondrial dysfunction and disrupted cellular energetics (Chung et al., 2019; Rojas-Morales et al., 2019). Whether targeted dietary manipulation of isoleucine can modulate these processes and improve cardiac outcomes remains unclear.

In this study, we sought to define the contribution of dietary isoleucine restriction (IleR) to cardiac adaptation under pressure overload and to delineate the underlying mechanisms. Building on prior observations that lowering total BCAA intake improves cardiac outcomes, we tested the hypothesis that isoleucine restriction is sufficient to confer cardioprotection by preserving mitochondrial function. Since IleR resulted in hypophagia (i.e., caloric restriction), we further examined the extent to which its effects overlap with those of calorie restriction and investigated the role of FOXO3-mediated transcriptional programs in maintaining mitochondrial gene expression. Understanding how specific dietary components regulate cardiac metabolism and function may provide new avenues for therapeutic intervention in heart failure.

## Results

### An isoleucine-restricted (IleR) diet improves the outcome of pressure overload-induced cardiac hypertrophy

We previously reported that dietary restriction of branched-chain amino acids (BCAAs) improves outcomes in pressure overload–induced cardiac hypertrophy (Yang et al., 2023). Further investigation identified isoleucine as the primary mediator of this effect. When mice that are maintained on a control or an isoleucine-restricted (IleR) diet were subjected to transverse aortic constriction (TAC), they exhibited equal increases in heart weight/body weight (HW/BW) after 1 week on either diet (1W, Fig. 1A). However, within 2W, the increase in HW/BW of mice on an IleR diet plateaued, while in mice on the control diet it continued to increase for up to 4W (Fig. 1B–C). The latter culminated in a reduced ejection fraction, which was preserved with the IleR diet (Fig. 1D). Note that the IleR diet is Ile-free but reduces circulating isoleucine levels by only ∼60% (supplementary Figure 1), as the gut microbiome supplies substantial levels of circulating BCAA (Pedersen et al., 2016).

**Figure 1.**
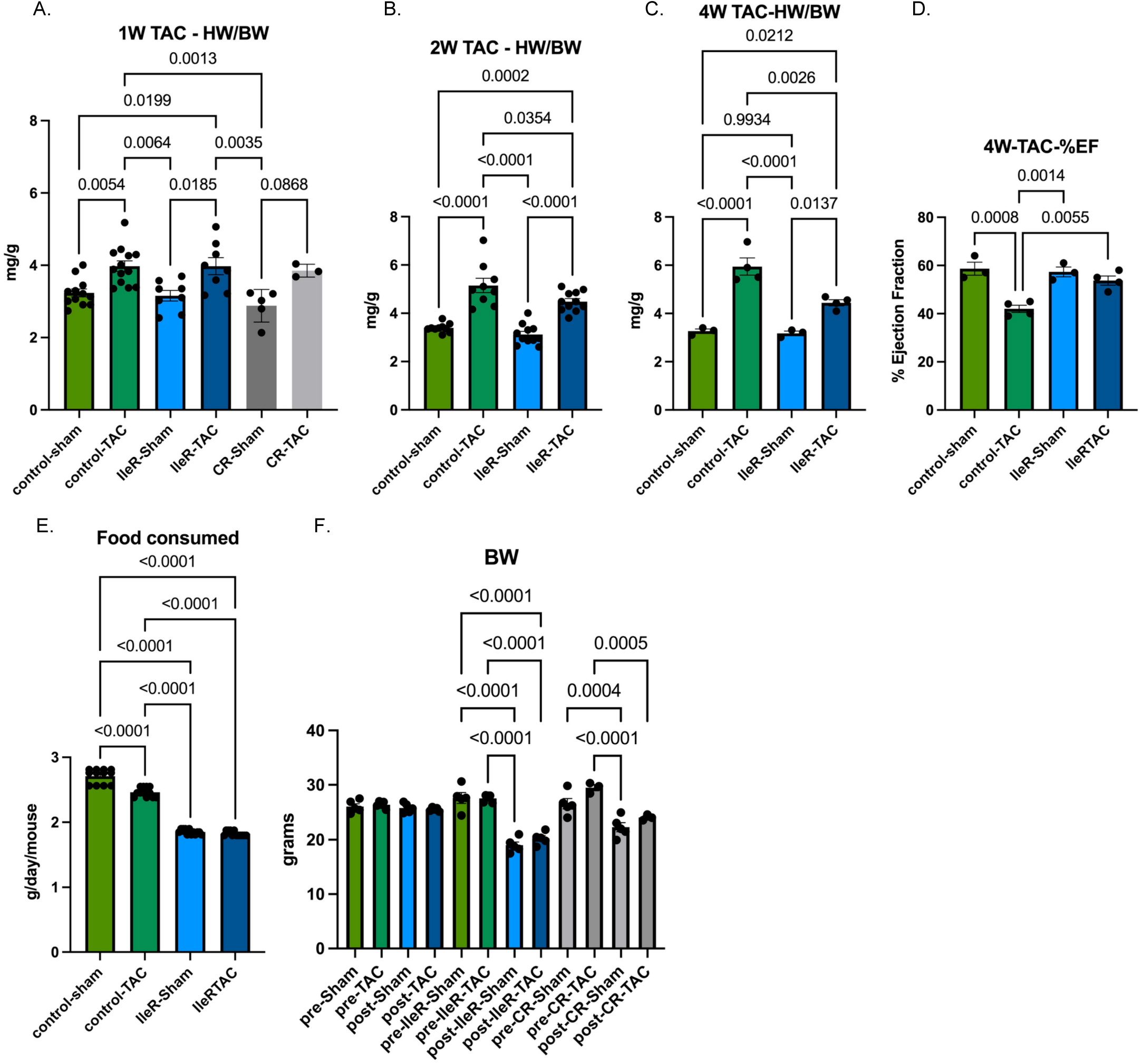
IleR decelerates the development of cardiac hypertrophy and failure. Mice were fed an IleR or CR diet 4 days before they were subjected to transverse aortic constriction (TAC) or a sham operation, then maintained on the same diets for **A.** one week (1W), B. 2W, or **C-D.** 4W, after which the hearts were analyzed by echocardiography (n=3-12). **A-C.** HW/BW (mg/g) and D. %EF after 4W of TAC were averaged and plotted. **E.** Food consumption was measured by weighing the amount of food before and after adding it to the mice cages, divided by the number of mice/cage/day. The grams consumed/day/mouse are graphed (15 mice, 5 mice/cage, each). **F.** Mice were weighed before and after, pre- and post-TAC or sham surgeries, while on a control, IleR, or CR diets for a total of 11 days (4 days before surgeries and 1 W after surgery). Mice on a CR diet were fed 1.8 g/mouse/day, equivalent to the amount consumed by the mice on an IleR diet. Body weight (grams) was averaged and graphed. **A-F.** Error bars represent SEM. The data were analyzed by one-way ANOVA; p-values <0.05 are indicated above the brackets.

The IleR diet triggered a reduction in food intake within 2 days of approximately 20–25% in mice, paralleled by a proportional decrease in body weight within 1W (Fig. 1E–F). When we fed the mice an equivalent calorie-restricted (CR) diet (1.8 g of control diet/mouse/day), it produced a similar reduction in body weight as did the IleR diet (Fig. 1E–F, gray bars). These mice also responded to pressure overload by an increase in HW/BW within 1W.

These findings suggest that isoleucine restriction tempers cardiac remodeling through both direct cardiac effects via histone propionylation, as we previously demonstrated (Yang et al., 2023), and indirect mechanisms related to reduced caloric intake. Although valine and leucine were similarly decreased, prior work demonstrates that only isoleucine specifically affects mitochondrial respiration (Yang et al., 2023).

### Isoleucine- and calorie-restricted diets prevent pressure overload-induced decline in the expression of mitochondrial electron transfer chain subunits and mitochondrial respiration

To contrast the effects of IleR and CR on gene expression in the heart before or after imposing pressure overload, we performed RNA-seq analysis of hearts from mice maintained on either diet, for 1W after a sham or transverse aortic constriction (TAC) surgery. Gene ontology analysis revealed that the top downregulated pathways after TAC in mice on the control diet (padj ≤ 0.05) were related to mitochondrial respiration and the electron transport chain (ETC). The data show that the transcription of all expressed ETC components (Complexes I–V) was significantly reduced in the stressed hearts as demonstrated in the heatmap in Figure 2 (log₂ TAC/Sham; Fig. 2A–B). This effect was not attributable to reduced mitochondrial abundance, as mitochondrial content and expression of membrane protein markers (Timm and Tomm) were unchanged (Fig. 2A–B). Importantly, the TAC-induced suppression of ETC gene expression was completely abolished by either an IleR (log₂ TAC^IleR^/Sham^IleR^) or CR (log₂ TAC^CR^/Sham^CR^) diet (Fig. 2A-B), suggesting a shared regulatory mechanism. Accordingly, the log_2_ TAC^IleR^/TAC or TAC^CR^/TAC ratios reflect relatively higher ETC expression in IleR and CR hearts after stress compared with those on the control diet. Timm and Tomm were not significantly changed and were included as internal controls. Only significantly altered genes (padj ≤ 0.05) were included in the analysis for CxI-CxV; Timm and Tomm served as internal controls that were not significantly altered.

**Figure. 2.**
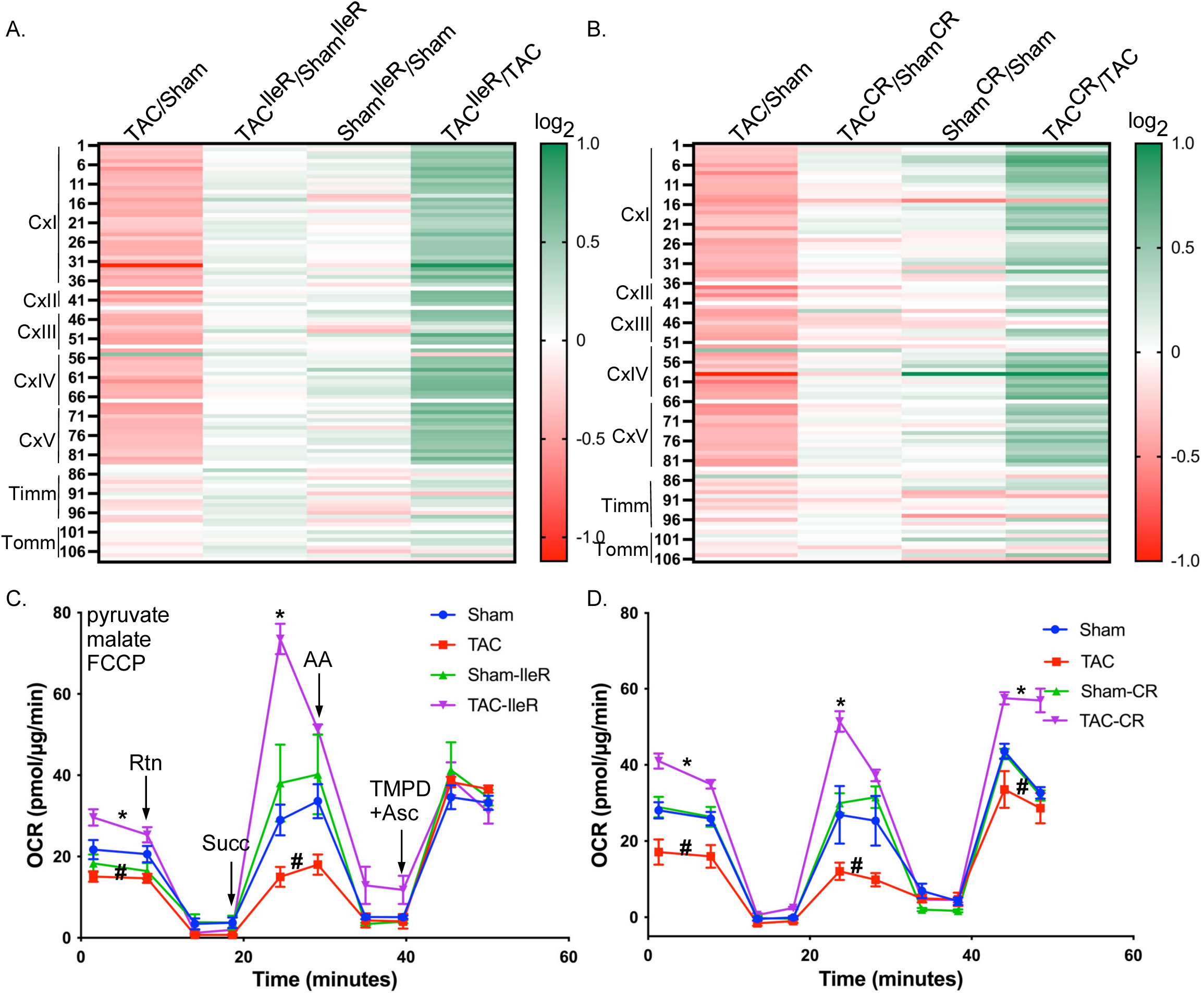
Cardiac pressure overload is associated with a decline in ETC (CxI-CxV) gene expression and electron flow, which is rescued by IleR or CR. Mice hearts were subjected to transverse aortic constriction (TAC) or a sham operation, then maintained on a **A.-B.** control, **A.** IleR, or **B.** CR diet for one week, after which the hearts were analyzed by RNA-Seq. The results for the genes of the ETC CxI-CxV are displayed in the heatmaps as the Log2 fold changes of **A.** TCA/Sham (control diet), TAC^IleR^/Sham^IleR^, Sham^IleR^/Sham, TAC^IleR^/TAC, and **B.** TCA/Sham (control diet), TAC^CR^/Sham^CR^, Sham^CR^/Sham, TAC^CR^/TAC. Only those with a significant change in TAC/Sham (control diet) were selected for the heatmap. **C-D.** Mitochondria were isolated from the hearts of mice similarly treated as described in **A-B**. and analyzed by an electron flow assay in the presence of pyruvate, malate, and FCCP, and monitored by sequential injections of rotenone (Rtn), succinate (Succ), antimycin A (AA), and tetramethylphenylenediamine (TMPD)+Asc (n=10). * p<0.05 v. Sham, # p<0.05 v. Sham^IleR^ or Sham^CR^, at the corresponding time points.

To determine functional consequences of the above results, we assessed mitochondrial electron flow in isolated cardiac mitochondria. Pressure overload reduced Complex I-III activities, whereas IleR significantly maintained these functions, increasing Complex I activity by 1.25-fold and Complex II by 2.3-fold (Fig. 2C). Similarly, CR-fed mice also showed improved respiration consistent with the gene expression results (Fig. 2D). In conclusion, IleR or CR diet preserves mitochondrial transcriptional and functional integrity during cardiac stress, with the most prominent activation being of Complex II, which supports improved respiratory capacity and metabolic adaptability under increased energetic demand (Pfleger et al., 2015). The results suggest that the IleR diet mediates its effect on mitochondrial respiration by reducing food intake, i.e., caloric restriction.

Mitochondrial respiration positively correlates with cristae abundance and morphology (Patten et al., 2014). To substantiate the observed changes in ETC gene expression and mitochondrial function, we performed electron microscopy on cardiac tissue from mice maintained on control or isoleucine-restricted (IleR) diets following 1W of sham or TAC surgery. In mice on the control diet, we observe 27±4 cristae/mitochondrion in the sham heart, which drops to 5±3 after 1W of pressure overload (Fig. 3A-B, E). In contrast, mice on an IleR diet did not exhibit a drop in cristae numbers, 30±8 and 28±5 cristae/mitochondrion in the sham and TAC heart, respectively (Fig. 3C-D, E). Additionally, these changes align with changes in the expression of Mitochondrial Contact Site and Cristae Organizing System (MICOS) genes, which regulate and maintain cristae formation (Fig. 3F). Collectively, these results demonstrate that isoleucine restriction preserves mitochondrial ultrastructure during cardiac stress, providing a structural basis for sustained respiratory capacity and metabolic resilience.

**Figure 3.**
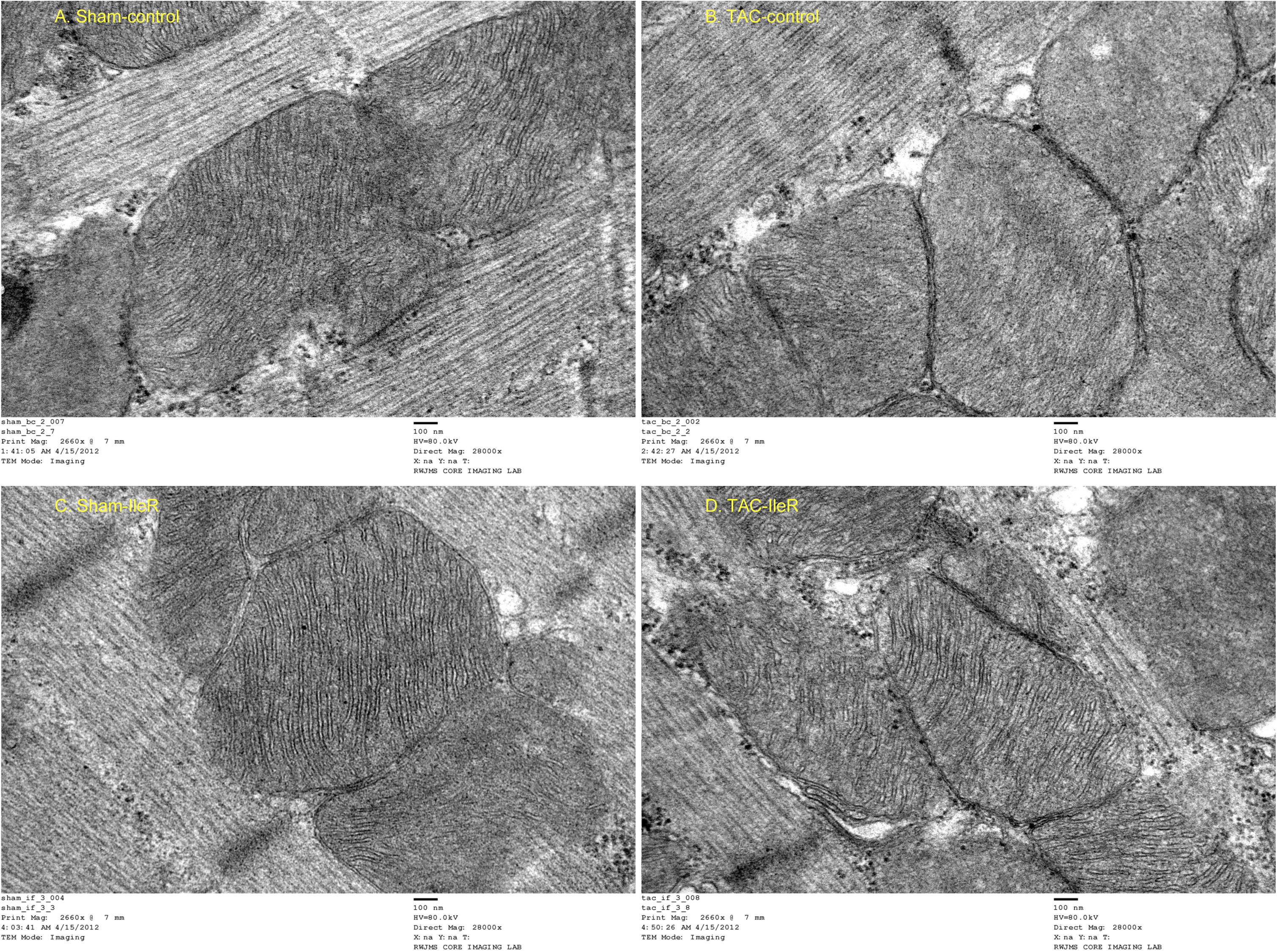
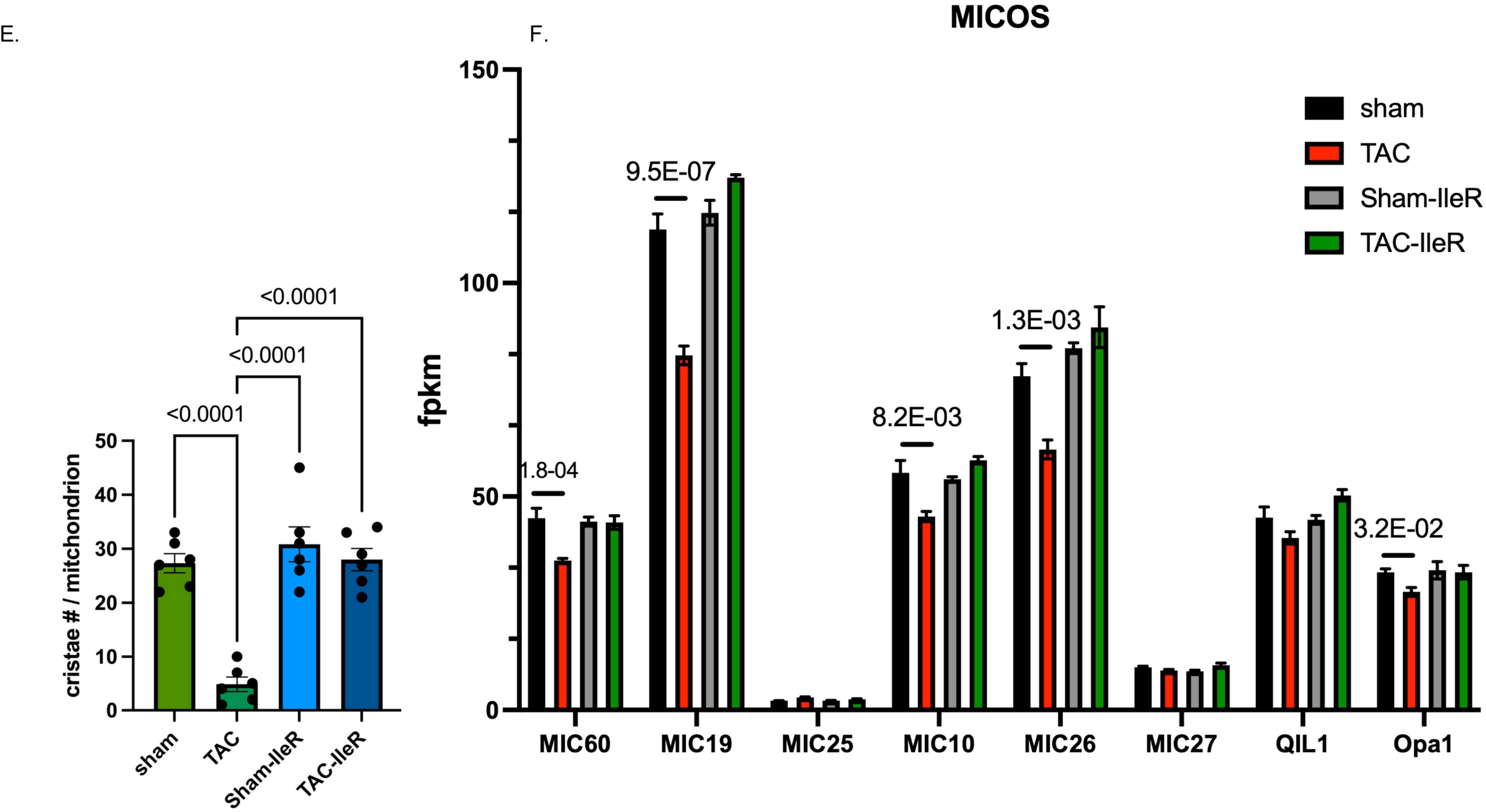
An isoleucine-restricted diet prevents dismantling of cristae during cardiac pressure overload. Mice were maintained on an **A.-B.** control or C.-D. IleR diet for 4 days before being subjected to a sham or TAC surgery. After 7 days the hearts were harvested and processed for EM, mag. 28000x (n=3, each). **E.** Cristae/mitochondrion were counted, averaged, and graphed (n=6 mitochondria). Values were analyzed by one-way ANOVA. P-values are shown on top. **F.** The RNA-Seq results (fpkm) for the MICOS (Mitochondrial Contact Site and Cristae Organizing System) subunits were graphed (n = 3). P-values on top of bars are pAdj for the RNA-Seq results.

### Isoleucine or glucose restriction directly enhances the respiratory capacity of rat cardiac myocytes and human iPSC-CM

To determine whether the mitochondrial protective effects of IleR or CR are direct or indirect on the heart, we examined isolated rat cardiac myocytes cultured with increasing concentrations of isoleucine (0–0.8 mM; standard concentration in media = 0.8 mM) or glucose 0-17.5 mM glucose (17.5 mM standard in DMEM/F12 and 5.5 mM standard in DMEM). After 24 h, oxygen consumption rate (OCR) was assessed using a mitochondrial stress test. The results show that a concentration of 0.08 mM Ile produced the highest basal (Fig. 4B), ATP-linked (Fig. 4C), and reserve respiratory capacity (RRC) OCR values, with the most pronounced effect on RRC (Fig. 4D). Likewise, OCR measurements after 24 h revealed maximal respiration with low glucose (2 mM), plateauing between 2–12 mM, with a marked decline at 17.5 mM, almost to levels comparable with glucose deprivation (Fig. 4E–H). These experiments were performed in the presence of 0.8 mM isoleucine (standard concentration in DMEM and most media), which, as seen in the previous experiment, imposes metabolic stress (i.e., suboptimal respiration). In conclusion, both isoleucine restriction and reduced glucose concentration directly enhance mitochondrial respiratory capacity in cardiomyocytes, supporting a cell-autonomous mechanism underlying the protective effects of IleR and CR.

**Figure 4.**
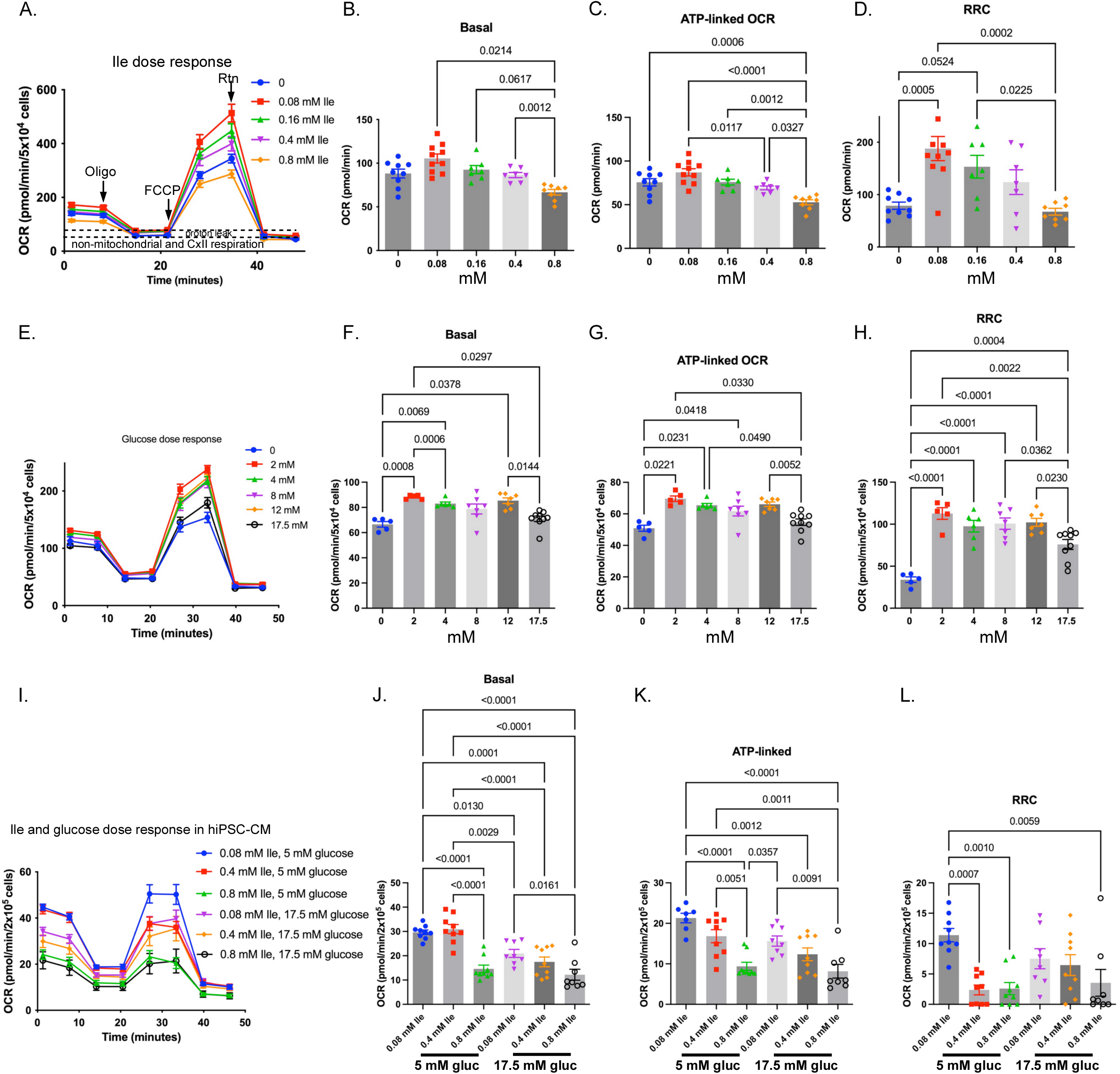
Reducing isoleucine (Ile) or glucose in cell culture medium increases mitochondrial respiration in cardiac myocytes. A-H. Neonatal rat cardiac myocytes were cultured in, **A-D.** custom-made DMEM without Ile, to which increasing doses of isoleucine were added (0-0.8 mM Ile, where 0.8 mM is the concentration found in conventional DMEM) for 24 h with no FBS, or **E-H.** in DMEM without glucose, to which increasing doses of glucose were added for 24 h with no FBS. **I-L.** Human iPCS-derived cardiac myocytes were treated with various doses of Ile and glucose. Cells were assayed by the Seahorse extracellular flux analyzer with the sequential addition of oligo, FCCP, and Rtn (n=7-10). **A, E, I.** The data were graphed as pmol/min/# of cells vs. time. Mitochondrial respiration (total OCR minus non-mitochondrial respiration) was calculated and graphed for **B, F, J.** basal, **C, G, K.** ATP-linked, and **D, H, L.** RRC OCR. Error bars represent SEM, and the data were analyzed by one-way ANOVA; p-values <0.05 are indicated above brackets.

To determine whether human iPSC-derived cardiomyocytes (iPSC-CMs) respond similarly to isoleucine restriction (IleR) or caloric restriction (CR), we cultured spontaneously beating cells in DMEM with varying concentrations of Ile (0.08, 0.4, or 0.8 mM) and glucose (5 or 17.5 mM). A mitochondrial stress test revealed that reducing isoleucine to 0.08 mM maximized basal, ATP-linked, and reserve respiratory capacity (RRC), with the most pronounced effect on RRC (Fig. 4I–L). Lower glucose levels (5 vs. 17.5 mM) further enhanced respiration. Conversely, high isoleucine (0.8 mM) consistently lowered OCR, irrespective of glucose concentration, whereas reducing isoleucine under high-glucose conditions markedly improved respiration. The highest OCR was observed with 0.08 mM Ile and 5 mM glucose, indicating synergy. In conclusion, IleR directly enhances mitochondrial respiratory capacity in rat and human cardiac myocytes, and its effects are modulated by glucose availability, supporting both conserved and context-dependent mechanisms across species and maturation states.

### Improved NAD⁺ homeostasis by IleR and CR diets during pressure overload on the heart

Maintaining the NAD^+^/NADH ratio is essential for sustaining the mitochondrial respiratory capacity. This ratio declines after 1W of pressure overload (Fig. 5A-C). RNA-seq analysis revealed that this is likely due to downregulation of genes involved in NAD⁺ biosynthesis via the salvage pathway, including *Nampt, Naprt,* and *Nmnat* (Supplementary Figure 2). In contrast, expression of NAD⁺-consuming enzymes, including *Cd38, Parps,* and tankyrase (*Tnks/Parp5*), was increased. Intriguingly, these changes were blocked by an IleR or CR diet (Fig. 5A-C and supplementary Figure 2). Increases in ATP (Fig. 5D) and Nampt protein (Fig. 5E-F) in the heart corroborate these results. The pressure overload-induced increase in Ankrd1, a marker of cardiac hypertrophy, did not differ across the diets and served as an indicator of the extent of hypertrophic growth.

**Figure 5.**
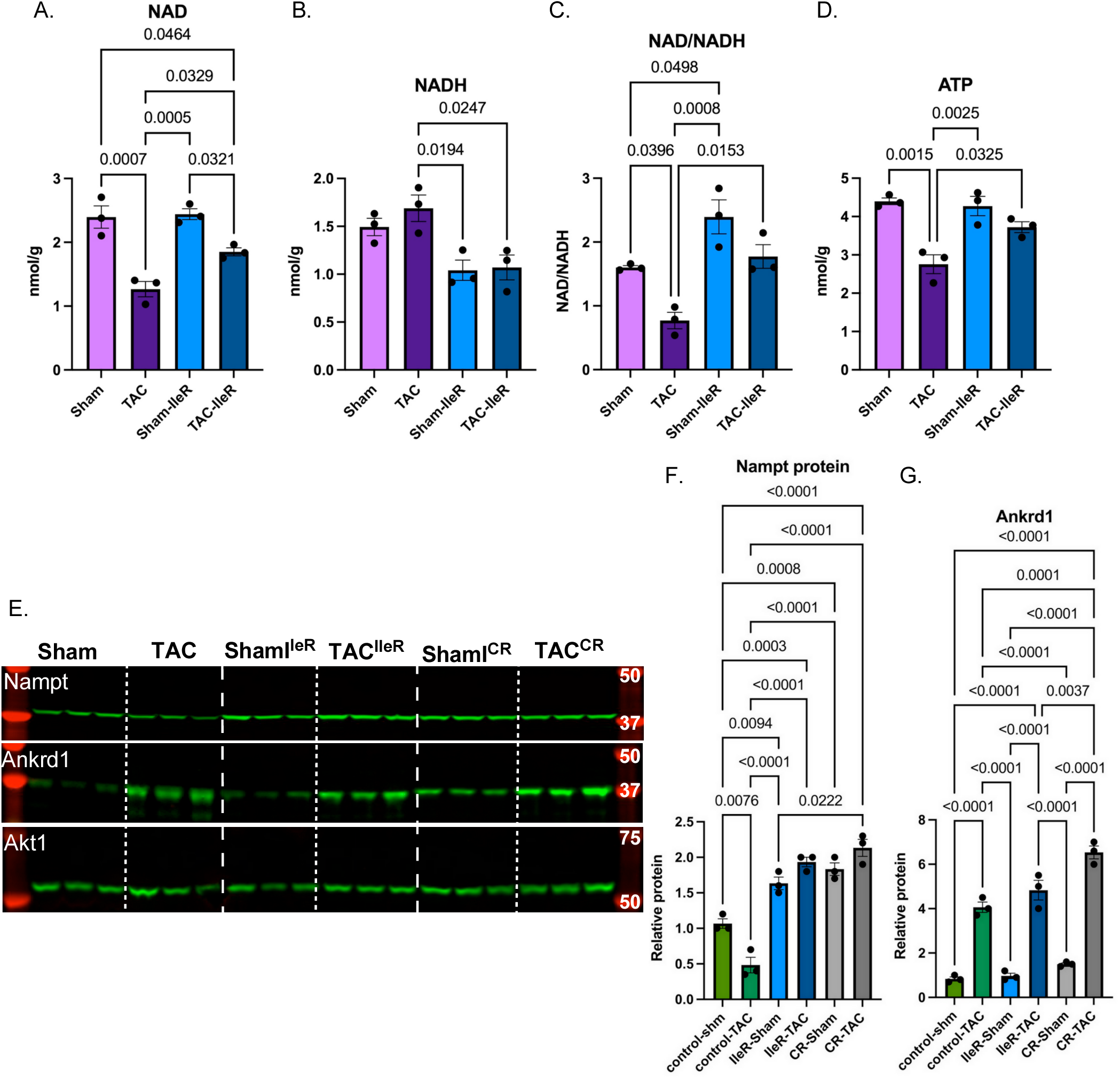
IleR maintains NAD+/NADH levels during pressure overload on the heart. Mice were fed a control or IleR diet for 4 days before subjecting them to a sham or TAC surgery. A-D. After one week, the hearts were extracted and analyzed by mass spectrometry for quantitation of NAD, NADH, and ATP (n=3, each). The results (nmol/g) were graphed, and the ratio of NAD/NADH was calculated. E. Protein extracted from the hearts of similarly treated mice was analyzed by WB. F-G. The signals for Nampt and Ankrd1 were quantitated, normalized, and graphed (n=3, each). Error bars represent SEM, and the data were analyzed by one-way ANOVA, and p-values <0.05 are listed on top.

To assess the functional impact of Nampt on mitochondrial respiration, cardiac myocytes were treated with increasing doses of either a Nampt activator (Nampt activator-1) or the inhibitor CHS-828 for 24 h. Nampt activation increased basal, ATP-linked, and RRC OCR in a dose-dependent manner, with maximal effects at 6 µM (Fig. 6A–D). Basal OCR increased by ∼35% (from 71±7 to 96±10 pmol/min), while RRC increased by ∼77% (from 117±30 to 208±72 pmol/min). Conversely, dose-dependent inhibition of Nampt reduced RRC by ∼60% (from 104±31 to 40±8 pmol/min) at 80 nM CHS-828 (Fig. 6E–H). Conversely, pharmacological inhibition of Cd38, a major NADase, using compound 78c, dose-dependently increased RRC by over twofold (Fig. 6I–L), with modest increases in basal and ATP-linked OCR. Collectively, these findings demonstrate that enhancing NAD⁺ biosynthesis or limiting NAD⁺ consumption improves mitochondrial reserve respiratory capacity, particularly RRC, in cardiac myocytes.

**Figure 6.**
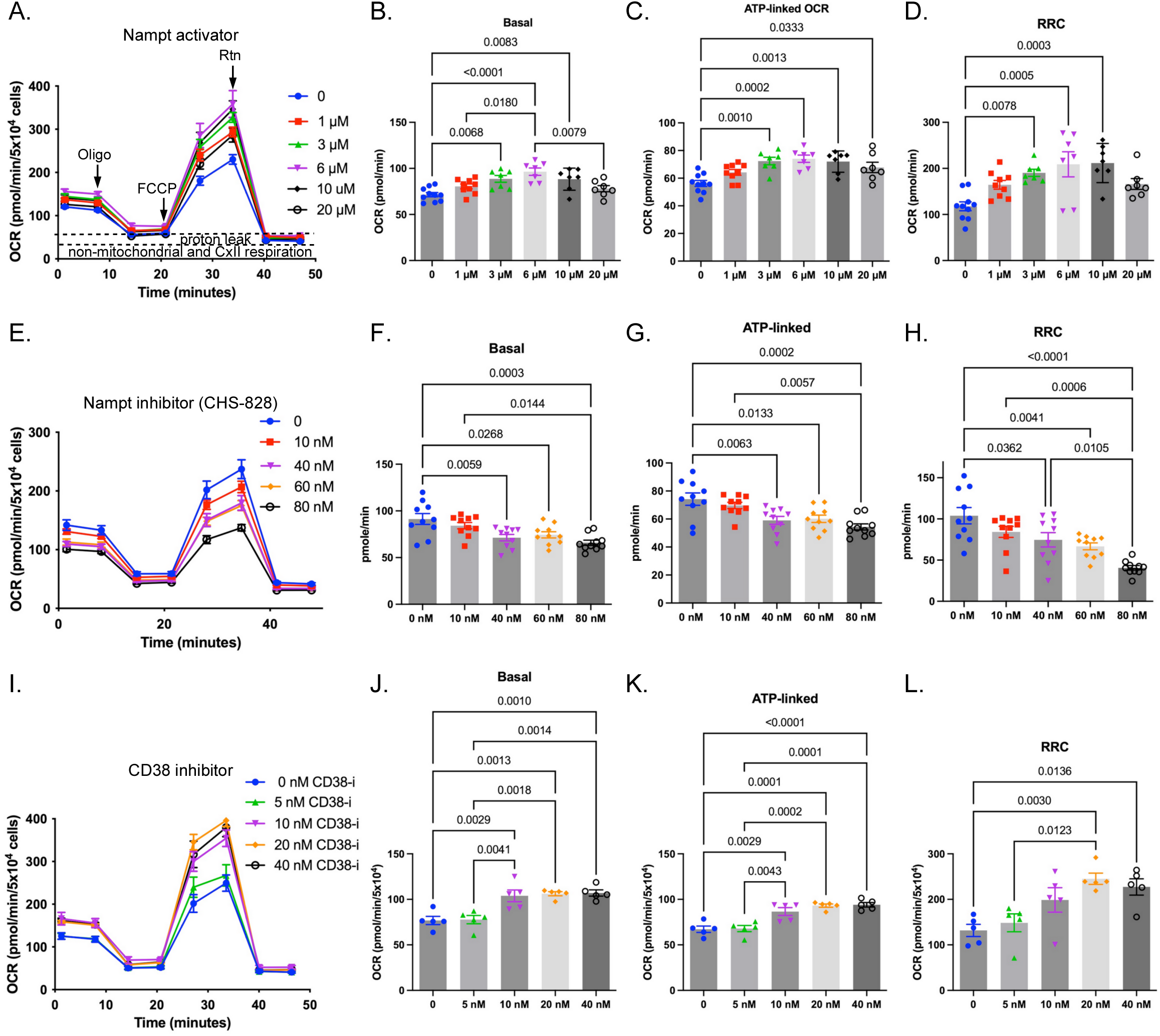
Activation of Nampt or inhibition of Cd38 enhances RRC in cardiac myocytes. A-D. Neonatal rat cardiac myocytes were incubated with increasing doses of the Nampt activator-1 (n=7-10), E-H. the Nampt inhibitor CHS-828, or I-L. Cd38 inhibitor CD38i (n=9-10). A, E. The data were graphed as pmol/min vs. time. Mitochondrial respiration (OCR minus non-mitochondrial respiration) was calculated and graphed for B, F, J. basal, C, G, K. ATP-linked, and D, H, L. RRC OCR. Error bars represent SEM, and the data were analyzed by one-way ANOVA; p-values for those <0.05 are indicated above brackets.

### Foxo3a and its targets are downregulated in the heart during pressure overload, which is abolished by IleR or CR diets

IleR and CR diets completely prevent the downregulation of ETC genes (Fig. 2), prompting us to explore the identity of the transcription factors responsible for this effect. Our data indicate that pressure overload reduces Foxo3 expression (Log_2_ TAC/Sham; Fig. 7-B), whereas IleR and CR increase Foxo3 levels in the normal/sham hearts (Log_2_ Sham^IleR^/Sham and Sham^CR^/Sham; Fig. 7A–B). This increase counteracts the pressure overload–mediated decrease (Log_2_ TAC^IleR^/Sham^IleR^ or TAC^CR^/Sham^CR^; Fig. 7A–B), resulting in a net increase in Foxo3 expression in TAC^IleR^ and TAC^CR^ compared to TAC controls (Log_2_ fold change: +1.18 and +0.95, respectively; Fig. 7A–B).

**Figure 7.**
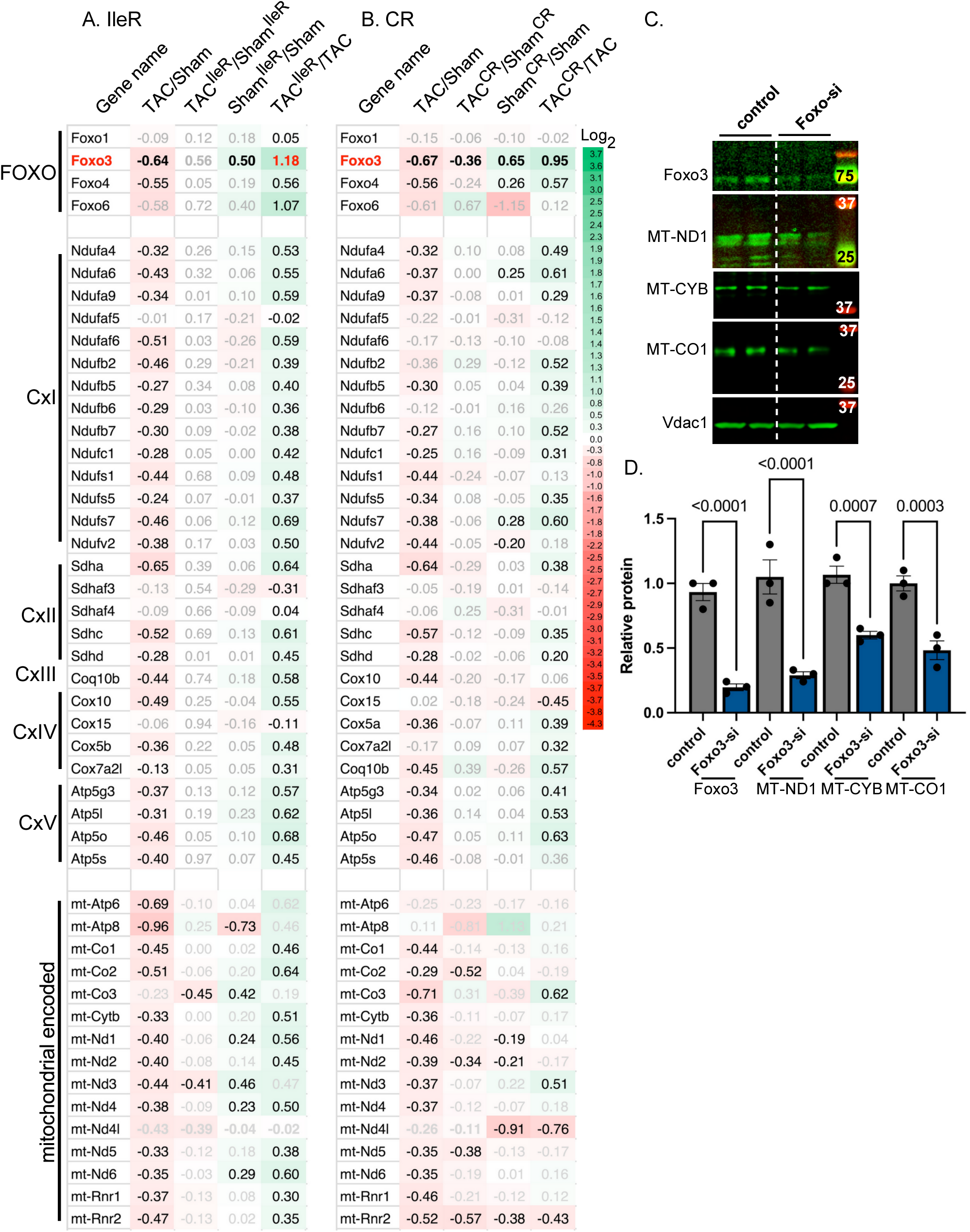
Foxo3 is upregulated by IleR and CR and increases the expression of mitochondrial genes. The hearts of mice fed an IleR or CR diet, subjected to sham or TAC surgeries, were analyzed by RNA-Seq. **A.-B.** The heatmap shows the differential RNA-Seq expression analysis (log2 fold change in fpkm of TAC vs. Sham) for Foxo3 and its ETC- and mitochondria-encoded gene targets. Numbers shown in black v. grey have a padj of ≤0.05 for the Log2 fold change of TAC/Sham, TAC^IleR^/Sham^IleR^, Sham^IleR^/Sham, TAC^IleR^/TAC, listed on top of each column. **C.** Neonatal cardiac myocytes were incubated with Ad5.short hairpin silencing RNA (Foxo-si) targeting Foxo3 for 24 h, after which mitochondria were isolated and analyzed by WB. **D.** The signals for Foxo3, MT-ND1, MT-CYB, MT-CO1 were quantitated, normalized to Vdac1, and graphed (n=3). Error bars represent SEM, and the data were analyzed by one-way ANOVA, and p-values <0.05 are listed on top.

Foxo3a transcriptional targets, identified in the ChEA dataset based on ChIP-chip, ChIP-seq, and related approaches (Diamant et al., 2025), include at least 28 ETC subunit genes and Opa1, a key regulator of cristae structure (Patten et al., 2014). Consistent with Foxo3 expression patterns, these targets are downregulated during pressure overload and restored by IleR or CR diets (Fig. 7A–B). Additionally, Foxo3 has been shown to bind mitochondrial gene regulatory regions, enhancing their expression and respiratory capacity (Peserico et al., 2013). Accordingly, IleR-and CR-driven Foxo3 upregulation is associated with preservation of mitochondrially encoded gene expression (Fig. 7A–B), further supported by reduced ND1, MT-CYB, and MT-CO1 expression following Foxo3 knockdown in cardiomyocytes (Fig. 7C).

Given Foxo3’s pro-apoptotic roles, we assessed caspase expression in response to pressure overload. Caspases displayed an inverse relationship with Foxo3 and ETC gene expression, increasing when these were decreased (Supplementary Figure 3). This induction was prevented by IleR and CR, consistent with improved mitochondrial function. In contrast, antioxidant and detoxification genes (e.g., Sod2 and Gstm family members) showed positive correlations with Foxo3 expression (Supplementary Figure 3). Collectively, these findings support a central role for Foxo3 in mediating the protective effects of IleR and CR against ETC gene downregulation.

Foxo3a is activated by dephosphorylation, which promotes its nuclear translocation and binding to target promoters. Western blot analysis confirmed that Foxo3a protein (72 kDa) is downregulated by pressure overload in the hearts of mice on a control diet; however, it increases in mice fed an IleR diet and is predominantly localized to the nucleoplasm, with partial association with chromatin (Fig. 8A–C). To assess its role in the mitochondrial respiratory capacity, it was knocked down in neonatal cardiac myocytes using Ad.Foxo3-shRNA (Foxo3-si). Foxo3 knockdown (by ∼70%, supplementary Figure 4) reduced basal and ATP-linked OCR by ∼ 50% while abolishing the mitochondrial RRC (Fig. 8D–G). These findings demonstrate that Foxo3 is essential for maintaining mitochondrial respiratory functions and mediates the beneficial effects of the IleR and CR diets on cardiac bioenergetics.

**Figure 8.**
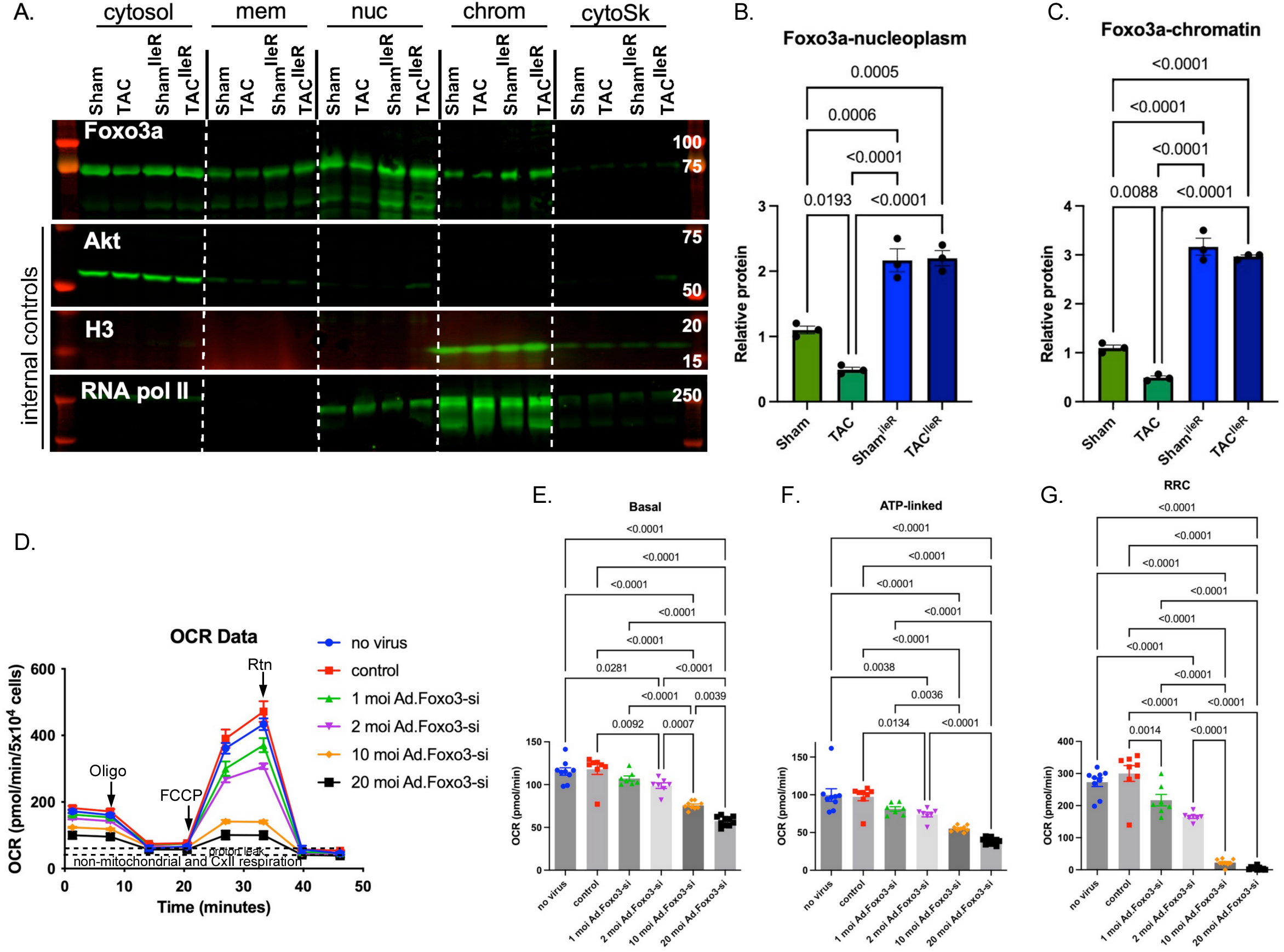
Foxo3a is required for mitochondrial respiration. **A**. Mice were fed an IleR diet 4 days before they were subjected to a sham or TAC surgery and maintained on the same diet for one week. Hearts were isolated, protein fractionated, and analyzed by WB. **B-C.** The WB signals were quantitated, normalized to internal control, averaged, graphed, and analyzed by one-way ANOVA (n=3). **D-G.** Neonatal cardiac myocytes were cultured with increasing doses of adenovirus harboring shRNA Foxo3 (Foxo3-si), or a control virus, for 24 h. Cells were assayed by the Seahorse extracellular flux analyzer with the sequential addition of oligo, FCCP, and Rtn (n=7-10). **D.** The data were graphed as pmol/min/cell number vs. real time. Mitochondrial respiration (total OCR minus non-mitochondrial respiration) was calculated and graphed for **E.** basal, **F.** ATP-linked, and **G.** RRC OCR. Error bars represent SEM, and the data were analyzed by one-way ANOVA; p-values <0.05 are indicated above brackets.

### Preservation of mitochondrial function is associated with reduced cardiac fibrosis in the stressed heart

The reduction in ETC gene expression, NAD^+^/NADH, and mitochondrial function during cardiac stress is associated with increased inflammation, cell death, and, consequently, increased fibrosis (Fig. 9C-D). Our RNA-seq analysis revealed a marked increase in collagen gene expression in the hypertrophied hearts, which was completely suppressed by IleR diet (Fig. 9A-B). The heatmap illustrates the log₂ fold changes in collagen isoforms, *Tgfb* and its receptors, and integrins across the various dietary conditions (log_2_ TAC/Sham control diet, and TAC^IleR^/Sham^Ile^ ^RIleR^ diet), as well as Sham^IleR^/Sham and TAC^IleR^/TAC comparisons (Fig. 9A). Comparatively, CR significantly reduced the basal levels of collagen gene expression, maintaining the fold increase induced by pressure overload, but with a net effect of overall a significant reduction collagen levels (Fig. 9B). The findings were validated by Picro Sirius Red staining (Fig. 9C–D) and Western blot analysis of Col1a1 (supplementary Fig. 6). These results corroborate the improved mitochondrial respiratory functions mediated by IleR and CR diets.

**Figure 9.**
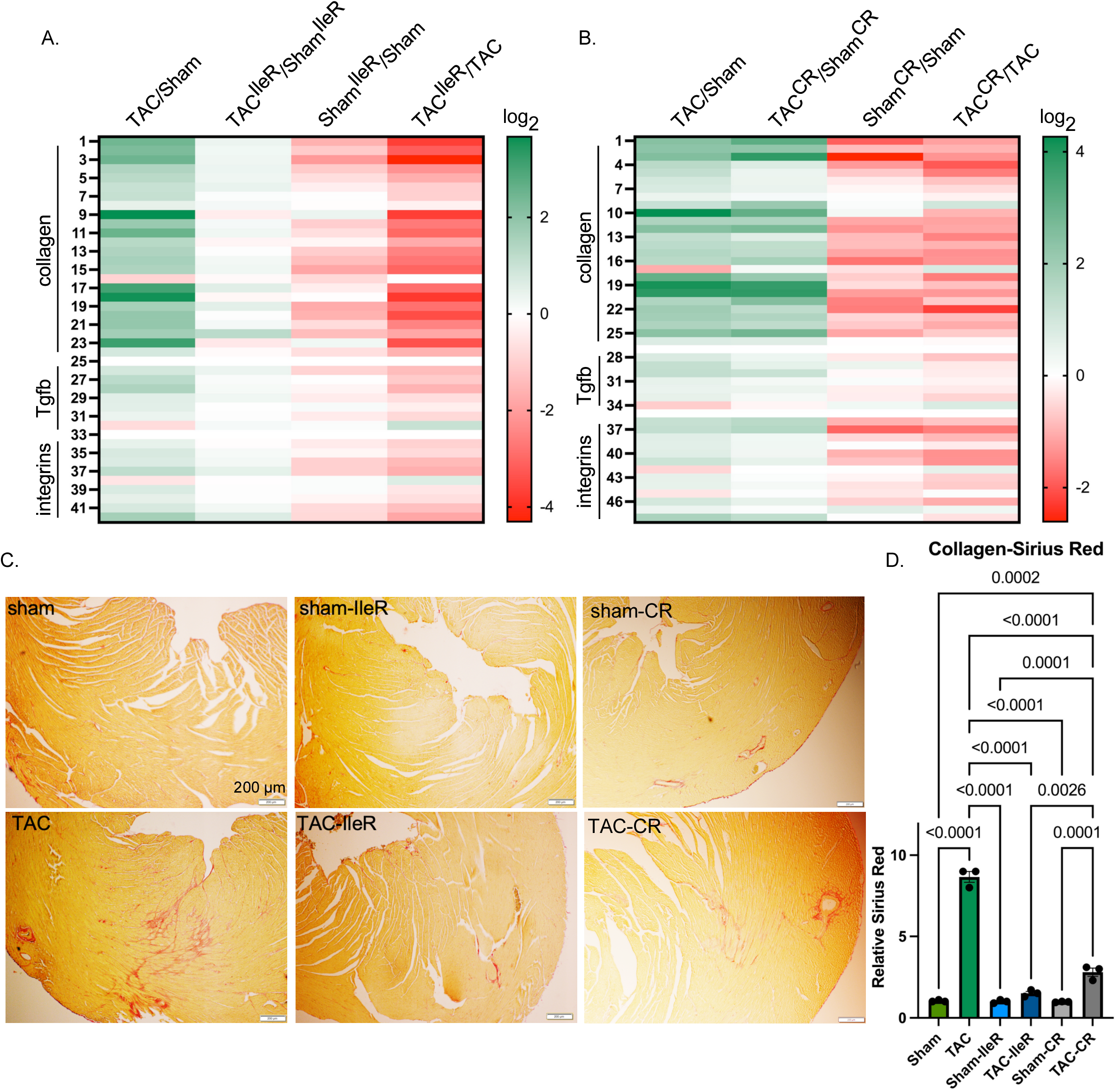
An IleR or CR diet reduces TAC-induced fibrosis. **A.** Mice were fed a control, IleR, or CR diet 4 days before they were subjected to a sham or TAC surgery and maintained on the same diet for one week. **A-B.** Total RNA was extracted from the heart and analyzed by RNA-Seq, in which collagen isoforms, TGFβ ligands and receptors, and integrins are displayed in the heatmaps as Log2 fold changes of **A.** TCA/Sham (control diet), TAC^IleR^/Sham^IleR^, Sham^IleR^/Sham, TAC^IleR^/TAC, and **B.** TCA/Sham (control diet), TAC^CR^/Sham^CR^, ShamCR/Sham, TAC^CR^/TAC. Only those with a significant change in TAC/Sham (control diet) were selected for the heatmap, or **C.** hearts were fixed and stained with Picro Sirius Red. **D.** Collagen (red) was quantitated, averaged, and graphed as relative values, after adjusting one of the sham values to 1 (n=3). The results were analyzed using one-way ANOVA, and p-values ≤0.05 are shown on the graph.

## Discussion

The present study identifies dietary isoleucine restriction (IleR) as a critical determinant of cardiac adaptation to stress/pressure overload and provides mechanistic insight linking amino acid availability, mitochondrial bioenergetics, and transcriptional control of metabolic remodeling. Building on our prior work implicating branched-chain amino acids (BCAAs) in cardiac pathology (Yang et al., 2023), we demonstrate here that isoleucine is the predominant BCAA mediating these effects. IleR attenuated the progression of pressure overload–induced hypertrophy, preserved systolic function, and improved mitochondrial integrity, supporting a model in which selective amino acid restriction reprograms cardiac metabolism to enhance stress resilience.

Previous work has shown that excess BCAA or isoleucine promotes hyperphagia (Liu et al., 2021; Yu et al., 2021). Conversely, we show that restricting dietary Ile reduces food intake and, in parallel, the mice’s weight by ∼20-25% within 4 days. The reduction was highly consistent between the animals (i.e., all responded) and plateaued thereafter. This suggests a highly conserved functional mechanism for Ile. Others and we have reported that Ile is responsible for histone propionylation and modulation of gene expression (Trefely et al., 2022; Yang et al., 2023). Additionally, Ile-induced hypophagia suggests that it may also mediate its beneficial effects via caloric restriction. Indeed, calorie restriction has been shown to confer health and lifespan benefits (Cohen et al., 2004; Lin et al., 2002; Nicolas et al., 1999; Wing et al., 1994). We tested this possibility by feeding mice a control diet with a caloric intake equivalent to that induced by the IleR diet and contrasted their effects on gene expression using RNA-Seq. As observed in Figure 2, the ETC gene expression profile was identical with both diets, as was the protection against mitochondrial respiratory decline induced by pressure overload. Both diets had similar effects on Foxo3 expression and mitochondrial-encoded gene expression, suppressing their decline during pressure overload (Fig. 7). Both diets also inhibited collagen expression in the heart, albeit with some minor differences. While the IleR diet completely inhibited pressure overload-induced upregulation of all collagen isoforms expressed in the heart, the CR diet exhibited similar increases in log_2_ fold change, but with overall significantly lower levels, as reflected in Figure 9 and Supplementary Figure 6 (log_2_ Sham^CR^/Sham and TAC^CR^/TAC). Other significant changes elicited by both diets include upregulation of several sarcomeric proteins (supplementary Figure 7) and suppression of histocompatibility genes (supplementary Figure 5), both at baseline and during cardiac stress, substantiating their overall beneficial effects on heart health, especially during stress.

Therefore, the question remains: are the effects of IleR all a consequence of calorie-restriction-induced metabolic remodeling? Measuring BCAA in the circulation before and after pressure overload, we find that isoleucine is reduced, as expected, with the IleR diet, in sham- and TAC-operated hearts (supplementary Figure 1). However, leucine and valine are also reduced, rendering it hard to dissociate their effects. With the CR diet, we also observe a reduction in isoleucine, especially after TAC, although it is not as pronounced as with the IleR diet (44% vs. 73% reduction) and occurs only after pressure overload. We addressed this issue by testing the effects of IleR and CR conditions on isolated cultured human and rat cardiac myocytes. Importantly, our in vitro studies establish that the effects of IleR on mitochondrial respiration are, at least in part, cell autonomous. Reducing isoleucine directly enhanced basal, ATP-linked, and reserve respiratory capacity (RRC) in both neonatal rat cardiomyocytes and human iPSC-derived cardiomyocytes within 24 h. The most robust effect was observed on RRC, indicating improved bioenergetic flexibility. Note, neither leucine nor valine had any effect on mitochondrial respiration (Yang et al., 2023). These findings align with broader literature demonstrating that nutrient restriction enhances mitochondrial efficiency and stress resistance (Broderick et al., 2002; Civitarese et al., 2007; Gredilla et al., 2001; Nisoli et al., 2005) and our previous report associating RRC with cell resistance to hypoxia and cell death (Pfleger et al., 2015). The interaction between isoleucine and glucose availability supports the concept that nutrient composition, rather than total caloric load alone, governs mitochondrial performance. Furthermore, these data suggest that Ile exerts both direct and indirect effects on cardiac myocytes. The former through modulating histone propionylation, as we have previously reported (Yang et al., 2023), and the latter through caloric restriction, plausibly through regulating satiety (Solon-Biet et al., 2019). Similarly, it has been reported that IleR (67% reduction) in aging mice promotes metabolic health in mice without reducing caloric intake (Yeh et al., 2024)

The preservation of mitochondrial function under IleR was further supported by ultrastructural analyses. Pressure overload disrupted cristae organization, whereas IleR maintained cristae density and morphology. Given that cristae structure is tightly coupled to respiratory efficiency (Patten et al., 2014), these findings provide a structural basis for maintained oxidative phosphorylation. Alterations in MICOS gene expression under IleR or CR suggest that regulation of cristae architecture contributes to this phenotype, in agreement with established roles of MICOS in maintaining inner mitochondrial membrane organization (van der Laan et al., 2016).

Mechanistically, we identify Foxo3 as a central regulator linking nutrient status to mitochondrial gene expression. Pressure overload suppressed Foxo3 expression, whereas IleR and CR enhanced it at baseline and prevented its decline during pressure overload. Foxo3 targets include numerous ETC subunits (harmonizome (Diamant et al., 2025)) and Opa1, a key regulator of mitochondrial dynamics and cristae structure (Patten et al., 2014). Restoration of these targets with IleR/CR diets aligns with preserved mitochondrial function and morphology. Prior studies have shown that Foxo3 transcription factor regulates oxidative metabolism and longevity (Chaanine et al., 2016; Morris et al., 2015; Shimokawa et al., 2015; Zhong et al., 2023) and can directly enhance the expression of mitochondrially encoded genes (Peserico et al., 2013). Loss- of-function experiments in our study confirm that Foxo3 is required for maintaining respiratory capacity, positioning Foxo3 as a key transcriptional effector of metabolic adaptation in the stressed heart.

The regulation of the mitochondrial respiratory function is multifactorial. It not only requires the maintenance of ETC subunit expression, but also its proper assembly on cristae, and the availability of the electron transporters, NAD^+^ and FAD. We find that all these parameters are reduced during pressure overload when the metabolic conditions are unfavorable, i.e., the diet includes relatively high levels of isoleucine and caloric intake (conventional chow: 10% Fat, 20% protein, and 70% carbohydrates). Pressure overload induces a transcriptional shift consistent with NAD⁺ depletion, characterized by reduced expression of biosynthetic enzymes and increased expression of NAD⁺-consuming enzymes. IleR and CR diets reversed these changes, restoring NAD⁺ homeostasis. Functional assays in our study demonstrated that enhancing NAD⁺ biosynthesis or inhibiting its consumption improves mitochondrial respiration, particularly RRC, in agreement with prior work on NAMPT (Oka et al., 2021) and CD38 modulation (Guan et al., 2016). Given that ETC expression and cristae formation are reduced by pressure overload, in addition to the reduction in NAD^+^/NADH, we conclude that simple replenishment of NAD^+^ or its precursors might not effectively improve disease conditions, as seen in some clinical trials (Poljšak et al., 2022). This underscores the effectiveness of modifying the diet for improving cardiovascular health and stress resistance.

Beyond bioenergetics, IleR diet exerted profound effects on pathological remodeling. Both transcriptomic and histological analyses revealed a marked reduction in cardiac fibrosis with IleR, with CR showing a similar but slightly less robust effect. These findings are consistent with studies linking improved mitochondrial function and NAD⁺ metabolism to reduced fibrosis and inflammation (Shi et al., 2021). Notably, the antifibrotic effect occurred independently of changes in hypertrophic markers such as Ankrd1, suggesting that metabolic remodeling can uncouple fibrosis from hypertrophic growth, reminiscent of physiological hypertrophy. Also, this highlights disruption of mitochondrial homeostasis as a key determinant of pathological remodeling independent of cardiomyocyte hypertrophy per se.

Several limitations should be acknowledged. First, although the dietary design isolates isoleucine deprivation, systemic metabolic adaptations—including contributions from the gut microbiome—may influence circulating amino acid levels. Second, while in vitro findings demonstrate direct cardiomyocyte effects, in vivo responses likely involve inter-organ metabolic crosstalk, particularly the liver. Third, additional signaling pathways, such as mTOR and integrated stress responses, may contribute to the observed phenotype and require further investigation. Finally, translation of isoleucine restriction to clinical settings will require careful evaluation of feasibility, safety, and long-term metabolic effects in humans.

In conclusion, this study demonstrates that selective restriction of isoleucine enhances mitochondrial respiratory capacity, preserves cardiac structure and function, and attenuates fibrosis during pressure overload. These effects are mediated through coordinated regulation of mitochondrial gene expression, cristae architecture, and NAD⁺ metabolism, with Foxo3 serving as a central transcriptional node. Indeed, both IleR and Foxo3 have been identified as longevity factors. Our findings highlight nutrient-specific modulation of cardiac metabolism as a promising therapeutic strategy and identify isoleucine restriction as a potential intervention to mitigate pathological cardiac remodeling.

## Materials and Methods

### Animals, diets, and animal care

Ten to 12-week-old, male, C57BL/6J mice were purchased from ‘The Jackson Laboratory’, as needed. Mouse diets were purchased from Research Diets, Inc., NJ, including a custom-made control diet (product # A12450K) and an Ile-free diet (product # A23012703); see supplementary Figure 8 data for detailed formulas. Sprague-Dawley dams with 1- to 2-day-old pups were purchased from Charles River Laboratories, MA. All animal procedures used in this study are in accordance with the US National Institute of Health *Guidelines for the Care and Use of Laboratory Animals (No. 85-23)*. All protocols were approved by the Institutional Animal Care and Use Committee at the Rutgers-New Jersey Medical School.

### Neonatal rat cardiac myocytes culture

Cardiac myocytes were cultured as described in our previous reports (Sayed et al., 2015). Briefly, hearts were isolated from 1-day-old Sprague-Dawley rats, and the cells were dissociated with collagenase. Cardiac myocytes were purified on a Percoll density gradient followed by differential pre-plating for 30 min, which depleted the non-myocytes. Myocytes were cultured in DMEM/F12 plus 10% fetal bovine serum (FBS).

### Human iPSC-derived cardiac myocytes

The cells were purchased from Fujifilm Cellular Dynamics catalog #01434 and cultured according to the supplier’s recommendation. Twenty-four hours before the planned experiment, the medium was changed to DMEM with various concentrations of isoleucine and glucose as indicated in Figure 4I-L.

### Mitochondrial electron flow assay and mitochondrial stress test

For the ‘mitochondrial electron flow assay’, mitochondria were isolated from the heart tissue using differential centrifugation in a buffer composed of 70 mM sucrose, 210 mM mannitol, 5.0 mM HEPES, 1.0 mM EGTA, and 0.5 % (w/v) fatty acid free BSA, pH 7. Using 40 µg of mitochondria, the assay was conducted in a buffer composed of 70 mM sucrose, 220 mM mannitol, 10 mM KH2PO4, 5 mM MgCl2, 2 mM HEPES, 1.0 mM EGTA, 0.2 % (w/v) fatty acid-free BSA, pH 7.2, with 10 mM pyruvate, 2 mM malate, and 4 µM FCCP. Using the Seahorse XF^e^96 Analyzer, basal oxygen consumption rates (OCR) were measured before and after sequential injections of 2 µM rotenone (Rtn), 10 mM succinate (succ), 4 µM antimycin A (AA), and 10 mM ascorbate (Asc) plus 100 µM TMPD, final concentrations.

The ‘mitochondrial stress assay’ was performed as we previously described (Jeon et al., 2021). Cardiac myocytes, Hap1 or Hap1ΔPCCA, cells were seeded in 96-well Seahorse analyzer plates (50,000 cells/well) in full medium - DMEM plus 10% FBS - overnight (∼20 h). The medium was then changed to DMEM with varying concentration of BCAA (from 0 to 2-fold the levels present in DMEM), with 17.5 mM glucose, or with 100 µM palmitate-bovine serum albumin (BSA), as indicated in the figures. Using the Seahorse XF^e^96 Analyzer, basal oxygen consumption rates (OCR) were measured in live cells, followed by measurements after the sequential addition of 1 µM oligomycin at 8 min of the start of measurements, 1 µM FCCP at 22 min, and 10 µM antimycin A plus rotenone at 35 min. Two readings were taken after each compound was injected and the results plotted in real-time in *p*mol/min vs. time (min) (Pfleger et al., 2015). The raw data were then exported to ‘Agilent Seahorse XF Mito Stress Test Report Generator’ file, for calculation and graphing of the mitochondrial basal, spare respiratory capacity, proton leak, and ATP production.

### Transverse aortic constriction (TAC) in mice

This was performed as described in our previous reports (Han et al., 2012; Sayed et al., 2007). Briefly, a 7-0 braided polyester suture was tied around the transverse thoracic aorta, against a 27-gauge needle, between the innominate artery and the left common carotid artery. Control mice were subjected to a sham operation involving the same procedure, minus the aortic constriction.

### Echocardiography and doppler measurements

This was performed as described in our previous reports (Han et al., 2012; Sayed et al., 2007). Briefly, transthoracic echocardiography was performed using the Vevo 3100 imaging system (Visual Sonics, Inc.) with a MX400-30MHz (mouse, cardiology) scanhead, encapsulated, transducer. Electrocardiographic electrodes were taped to the four paws, then one dimensional (1D) M-mode and 2D B-mode tracings were recorded from the parasternal short-axis view at the mid papillary muscle level. In addition, pulse-wave Doppler was used to measure blood flow velocity and peak gradient pressure in the aorta. For analysis, we used the Vevo 3100 Software (Vevo Lab v3.2.6), which includes: analytic software package for B-Mode (2D) image capture and analysis; cine loop image capture, display, and review; software analytics for advanced measurements and annotations; and physiological data on-screen trace.

### Subcellular fractionation of protein and Western blotting

Cellular protein (25-50 µg) was fractionated using the subcellular protein fractionation kit (Thermo Fisher, cat # 78840), according to the manufacturer’s protocols. The cellular fractions were separated on a 4% to 12% gradient SDS-PAGE (Criterion gels, Bio-Rad) and transferred to a nitrocellulose membrane. The antibodies used include AKT1 (Millipore, 07-416), Ankrd1, Foxo3a (Cell Signaling Technology, 2497T), Nampt (Abcam, ABE236874), Mt-ND1 (Abcam ab181848), Mt-Co1 (Cell Signaling Technology 55159T), Mt-Cyb (Cell Signaling Technology 54618T), VDAC1 (Genscript, cat # A01419), RNA pol II (Active Motif, cat # 102660). The Western blot signals were detected by the Odyssey imaging system (LI-COR) and quantitated using Image J.

### Electron microscopy

Hearts were fixed in 2.5% Glutaraldehyde + 4% Paraformaldehyde in 0.1M Cacodylate buffer pH 7.4. Tissue electron microscopy was performed using the Philips CM12 electron microscope with AMT-XR11 digital camera at Rutgers Robert Wood Johnson Medical School.

### RNA-Seq

Total RNA was extracted from heart tissue using the Trizol Reagent (Invitrogen). Library preparation of pol(A) mRNA and sequencing on an Illumina NovaSeq X Plus PE150 platform using a paired-end 150 bp sequencing strategy was performed by Novogene Corporation Inc.

FeatureCounts (2.0.6) was used to count reads mapped to each gene. FPKM (Fragments Per Kilobase of transcript sequence per Million base pairs sequenced) for each gene was calculated based on the gene length and the read count mapped to that gene, while accounting for sequencing depth. The DESeq2 R package (1.42.0) was used for differential expression. P-value is adjusted using Benjamini and Hochberg’s methods to control the error discovery rate (padj). The threshold of significant differential expression is padj ≤ 0.05.

### Mass spectrometry measurements of NAD, NADH, ATP, and amino acids

Heart tissue samples were analyzed by Creative Proteomics, NY. NAD, NADH, and ATP were measured in heart tissue using LC-MRM/MS on two LC-MS/MS systems. One set was run with positive-ion detection on an Agilent 1290 UHPLC system coupled to an Agilent 6495C QQQ mass spectrometer, with 0.1% formic acid in water and 0.1% formic acid in methanol as the mobile phase for binary solvent gradient elution. The second set was run on a Waters Acquity UPLC system coupled to a Sciex QTRAP 6500 Plus mass spectrometer with negative-ion detection. A tributylamine solution and acetonitrile were used as the binary solvent gradient eluent. The concentrations of the molecules were calculated by interpolating the linear regression curves for individual compounds, using internal standard calibration.

Plasma was analyzed by Creative Proteomics, NY. Free amino acids were measured in the plasma using an LC-MS based on MRM. The Shimadzu UPLC Nexera II interfaced with a Sciex QTRAP 6500+ mass spectrometer equipped with a TurboIonSpray (TIS) electrospray ion source. The column used was ACCQ-TAG Ultra C18 1.7μm, 2.1*100 mm (Waters) and was run at a flow rate of at 0.4 mL/min at 45°C. The gradient of the mobile phases A (0.1% formic acid in water) and B (0.1% formic acid in acetonitrile) was as follows: 4% B to 30% B in 7 min, then ramped up to 90% B in 0.5 min, held at 90% for 2.5 min, back down to 4% B in 0.5 min.

### Statistical analysis

One-way ANOVA with post hoc Tukey or multiple-comparison analysis, or one-way ANOVA with Šídák’s multiple-comparison test, was used to determine significance between more than 2 groups. T-test (equal variance, 2-tailed) was used to calculate the significance between 2 groups; p ≤ 0.05 is considered significant. For the RNA-Seq data, differential gene expression (log2 fold change, LFC) was detected by DESeq2 R package (1.42.0). P-value was adjusted using Benjamini and Hochberg’s methods to control the error discovery rate (padj). The threshold of significant differential expression is padj ≤ 0.05.

### Study approval

All animal procedures used in this study are in accordance with US National Institute of Health *Guidelines for the Care and Use of Laboratory Animals (No. 85-23)*. All protocols were approved by the Institutional Animal Care and Use Committee at the Rutgers-New Jersey Medical School.

## Data Availability

All the raw and processed RNA-Seq will be deposited in NCBI’s Gene Expression Omnibus (GEO). Submission will be completed before the manuscript is published.

## Acknowledgment

We thank Dr. Junichi Sadoshima, Chairman of the Department of Cell Biology and Molecular Medicine at Rutgers University, for his support and advice, and the National Institute of Health funding of grants R01 HL146537 and R01 HL157739 to the corresponding author, R01HL150059 to Danish Sayed, and S10OD025238 instrumentation grant for the Vevo 3100 echocardiography machine to the Department of Cell Biology and Molecular Medicine at Rutgers University.

