## Supplemental figures for "Restricting Dietary Isoleucine Promotes Foxo3-Dependent Mitochondrial Respiration by Reducing Caloric Intake"

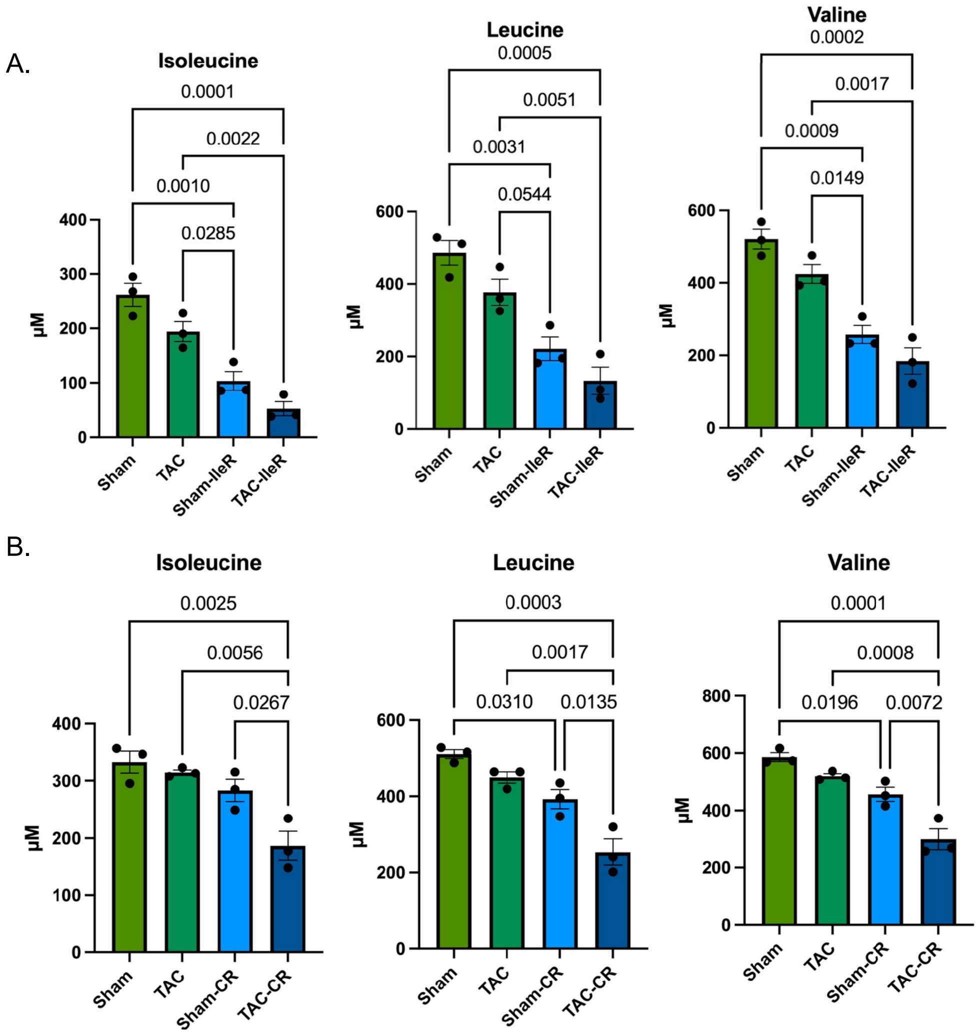

**Supplementary Figure 1. The effect of IleR and CR diets on plasma levels of essential amino acids.** Mice were fed a **A.** control or **A.** an isoleucine-free (IleR), or **B.** calorie-restricted (CR) diet 4 days before surgery and 7 days after a sham or TAC surgery. Plasma was then collected and amino acids were measured using mass spectrometry (n=3). **A-B.** Shows the BCAA. **C**. Shows all the essential amino acid after an IleR diet, and **D.** shows them after a CR diet. The results of plasma isoleucine, leucine, and valine concentrations (µM) were graphed, error bars represent SEM, the data were analyzed by one-way ANOVA; p-values for those <0.05 are indicated above the brackets.

C.

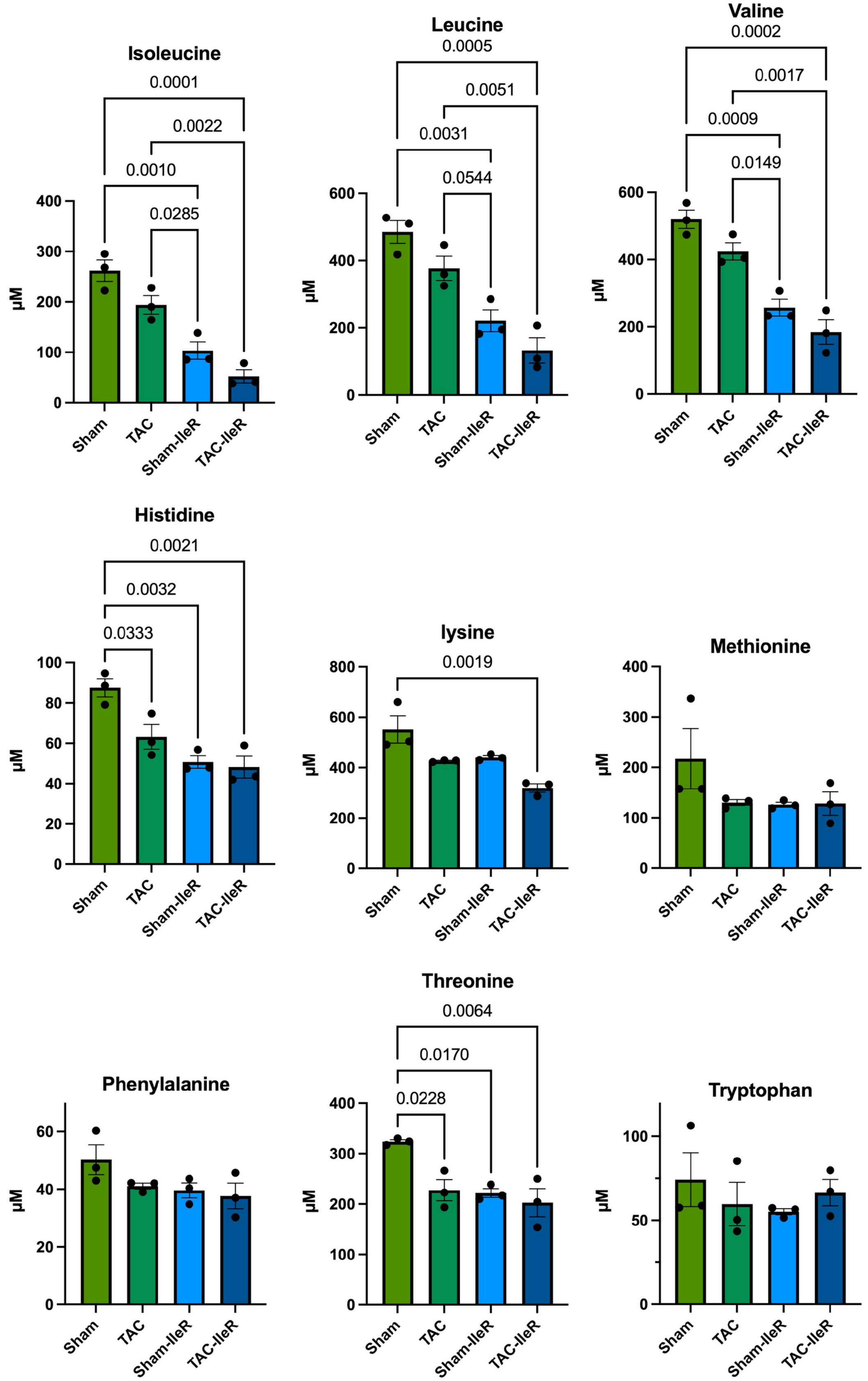

**Supplementary Figure 1 (continued). The effect of IleR and CR diets on plasma levels of essential amino acids. C**. Shows all plasma essential amino acid concnetrations after an IleR diet

D.

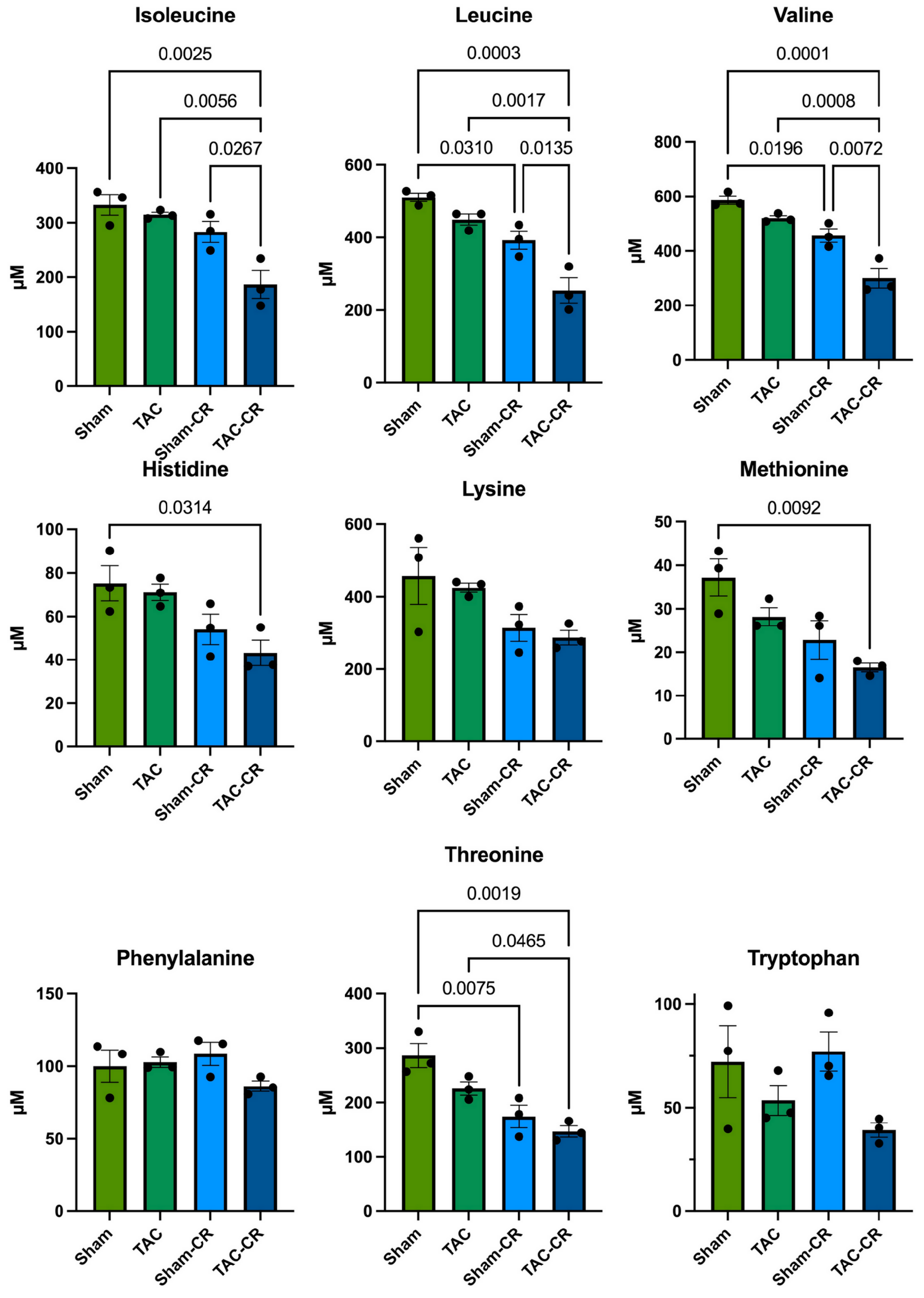

**Supplementary Figure 1 (continued). The effect of IleR and CR diets on plasma levels of essential amino acids. D**. Shows all plasma essential amino acid concnetrations after a CR diet

**A. IleR**

| | TAC/Sham | TAC$^{IleR}$/Sham$^{IleR}$ | Sham$^{IleR}$/Sham | TAC$^{IleR}$/TAC |
|---|---|---|---|---|
| Nadk2 | -0.92 | -0.24 | 0.36 | 1.02 |
| Nampt | -0.79 | -0.21 | 0.50 | 1.07 |
| Prkag1 | -0.58 | -0.14 | 0.33 | 0.75 |
| Nmnat3 | -0.57 | 0.15 | 0.10 | 0.80 |
| Sirt1 | -0.56 | -0.19 | 0.49 | 0.84 |
| Sirt3 | -0.55 | 0.05 | -0.24 | 0.34 |
| Prkab1 | -0.51 | -0.07 | 0.39 | 0.82 |
| Flad1 | -0.33 | 0.07 | -0.13 | 0.25 |
| Nmnat1 | -0.32 | 0.08 | -0.33 | 0.06 |
| Naprt | -0.30 | 0.15 | -0.41 | 0.02 |
| Parp1 | -0.25 | -0.09 | -0.16 | -0.01 |
| Prkaa2 | -0.18 | -0.06 | -0.07 | 0.04 |
| Parp4 | -0.10 | -0.09 | 0.30 | 0.28 |
| Parp2 | -0.02 | -0.16 | 0.17 | 0.02 |
| Parp6 | 0.02 | -0.18 | -0.11 | -0.32 |
| Prkaa1 | 0.06 | 0.04 | 0.15 | 0.11 |
| Nadk | 0.07 | -0.06 | -0.28 | -0.43 |
| Tnks2 | 0.10 | 0.03 | -0.24 | -0.33 |
| Parp8 | 0.12 | -0.16 | 0.24 | -0.05 |
| Parp11 | 0.13 | -0.08 | -0.27 | -0.49 |
| Prkag2 | 0.14 | 0.12 | -0.27 | -0.31 |
| Prkab2 | 0.21 | 0.10 | -0.19 | -0.31 |
| Tnks | 0.25 | 0.03 | 0.07 | -0.17 |
| Nadsyn1 | 0.28 | -0.1 | 0.23 | -0.2 |
| Tnks1bp1 | 0.40 | -0.11 | -0.19 | -0.71 |
| Parp12 | 0.44 | -0.12 | -0.11 | -0.68 |
| Parp3 | 0.49 | 0.29 | -0.16 | -0.38 |
| Cd38 | 0.53 | 0.24 | -0.70 | -1.00 |
| Parp14 | 0.59 | -0.16 | -0.51 | -1.28 |
| Parp10 | 0.64 | -0.34 | -0.27 | -1.27 |
| Parp9 | 0.79 | 0.04 | -0.61 | -1.38 |
| Parp16 | 0.81 | 0.10 | -0.32 | -1.04 |
| Nnmt | 1.14 | 0.33 | -0.73 | -1.56 |
| Nmrk2 | 2.09 | 0.59 | 0.24 | -1.27 |

**B. CR**

| | TAC/Sham | TAC$^{CR}$/Sham$^{CR}$ | Sham$^{CR}$/Sham | TAC$^{CR}$/TAC |
|---|---|---|---|---|
| Nadk2 | -0.98 | -0.47 | 0.59 | 0.59 |
| Nampt | -0.80 | -0.32 | 0.41 | 0.88 |
| Prkag1 | -0.47 | -0.03 | 0.20 | 0.63 |
| Nmnat3 | -0.69 | -0.15 | 0.18 | 0.71 |
| Sirt1 | -0.55 | 0.20 | 0.05 | 0.79 |
| Sirt3 | -0.29 | 0.02 | -0.19 | 0.11 |
| Prkab1 | -0.46 | -0.04 | 0.11 | 0.52 |
| Flad1 | -0.30 | -0.09 | -0.06 | 0.14 |
| Nmnat1 | -0.27 | -0.04 | -0.50 | -0.29 |
| Naprt | -0.32 | -0.11 | 0.13 | 0.33 |
| Parp1 | -0.29 | -0.19 | -0.11 | -0.02 |
| Prkaa2 | -0.22 | -0.23 | -0.23 | -0.24 |
| Parp4 | -0.08 | -0.43 | 0.49 | 0.13 |
| Parp2 | -0.12 | -0.11 | 0.08 | 0.08 |
| Parp6 | -0.24 | -0.03 | -0.24 | -0.03 |
| Prkaa1 | 0.11 | -0.05 | 0.04 | -0.13 |
| Nadk | 0.20 | -0.08 | -0.13 | -0.42 |
| Tnks2 | -0.03 | -0.07 | -0.32 | -0.37 |
| Parp8 | -0.07 | 0.09 | 0.09 | 0.25 |
| Parp11 | 0.13 | -0.12 | -0.14 | -0.40 |
| Prkag2 | 0.10 | 0.25 | 0.00 | 0.14 |
| Prkab2 | 0.21 | 0.23 | 0.23 | 0.24 |
| Tnks | 0.24 | 0.02 | -0.04 | -0.27 |
| Nadsyn1 | 0.09 | -0.1 | 0.28 | 0.05 |
| Tnks1bp1 | 0.48 | -0.10 | 0.28 | -0.31 |
| Parp12 | 0.36 | 0.04 | -0.19 | -0.52 |
| Parp3 | 0.40 | 0.36 | -0.19 | -0.24 |
| Cd38 | 0.61 | 0.30 | -0.65 | -0.96 |
| Parp14 | 0.67 | -0.15 | -0.28 | -1.11 |
| Parp10 | 0.78 | 0.11 | 0.05 | -0.63 |
| Parp9 | 0.82 | -0.30 | -0.07 | -1.20 |
| Parp16 | 0.66 | 0.44 | -0.15 | -0.38 |
| Nnmt | 1.01 | 0.26 | -0.03 | -0.79 |
| Nmrk2 | 2.04 | 1.09 | 0.18 | -0.77 |

**$Log_2$**

| $Log_2$ |
|---|
| 2 |
| 1.5 |
| 1.2 |
| 1 |
| 0.8 |
| 0.6 |
| 0.4 |
| 0.2 |
| 0 |
| -0.2 |
| -0.4 |
| -0.6 |
| -0.8 |
| -1 |
| -1.2 |
| -1.5 |

**Supplementary Figure 2. The decline in the NAD synthesis and the increase in NAD consuming enzymes in the heart during pressure overload and their rescue by IleR and CR diets.** Mice were fed a **A.-B.** control, **A.** an isoleucine-free (IleR), or **B.** calorie-restricted (CR) diet, 4 days before surgery and 7 days after a sham or TAC surgery. and maintained on the same diets for one-week, after which the hearts were analyzed by RNA-Seq. The results for the genes involved in NAD metabolism are displayed in the heatmaps as the $Log_2$ fold changes of **A.** TCA/Sham (control diet), TAC$^{IleR}$/Sham$^{IleR}$, Sham$^{IleR}$/Sham, TAC$^{IleR}$/TAC, and **B.** TCA/Sham (control diet), TAC$^{CR}$/Sham$^{CR}$, Sham$^{CR}$/Sham, TAC$^{CR}$/TAC. Only the significant (pAdj <0.05) $log_2$ values are displayed in black.

A. IleR

| Group | Gene name | TAC/Sham | $TAC^{IleR}/Sham^{IleR}$ | $Sham^{IleR}/Sham$ | $TAC^{IleR}/TAC$ |
|---|---|---|---|---|---|
| FOXO | Foxo1 | -0.09 | 0.12 | 0.18 | 0.05 |
| FOXO | **Foxo3** | **-0.64** | **0.56** | **0.50** | **1.18** |
| FOXO | Foxo4 | -0.55 | 0.05 | 0.19 | 0.56 |
| FOXO | Foxo6 | -0.58 | 0.72 | 0.40 | 1.07 |
| detoxification | Sod2 | -0.53 | 0.26 | -0.14 | 0.45 |
| detoxification | Gstm1 | -0.77 | 0.02 | 0.84 | 1.42 |
| detoxification | Gstm2 | -0.47 | 0.07 | 0.76 | 1.05 |
| detoxification | **Gstm4** | -0.37 | 0.14 | -0.23 | 0.38 |
| detoxification | Gstm5 | -0.51 | 0.46 | 0.17 | 0.58 |
| detoxification | Gstm7 | -0.73 | 0.22 | 0.88 | 1.47 |
| caspases | Casp12 | 0.79 | 0.21 | -0.44 | -0.94 |
| caspases | Casp2 | 0.31 | 0.65 | 0.11 | -0.14 |
| caspases | Casp3 | 1.63 | 0.96 | -0.32 | -1.96 |
| caspases | Casp4 | 0.65 | 0.61 | -1.38 | -1.79 |
| caspases | Casp6 | -0.10 | 0.90 | -0.41 | -0.31 |
| caspases | Casp7 | 0.66 | 0.27 | -0.26 | -0.71 |
| caspases | Casp8 | 0.95 | 0.83 | -0.52 | -1.44 |
| caspases | Casp9 | 0.12 | 0.29 | 0.05 | 0.09 |
| | Acta1 | 3.84 | 0.00 | -0.83 | -2.80 |
| | Myh6 | -0.43 | 0.67 | 0.15 | 0.60 |

B. CR

| Group | Gene name | TAC/Sham | $TAC^{CR}/Sham^{CR}$ | $Sham^{CR}/Sham$ | $TAC^{CR}/TAC$ |
|---|---|---|---|---|---|
| FOXO | Foxo1 | -0.15 | -0.06 | -0.10 | -0.02 |
| FOXO | **Foxo3** | **-0.67** | **-0.36** | **0.65** | **0.95** |
| FOXO | Foxo4 | -0.56 | -0.24 | 0.26 | 0.57 |
| FOXO | Foxo6 | -0.61 | 0.67 | -1.15 | 0.12 |
| detoxification | Sod2 | -0.53 | -0.19 | -0.29 | 0.04 |
| detoxification | Gstm1 | -0.82 | -0.31 | 0.68 | 1.19 |
| detoxification | Gstm2 | -0.54 | -0.07 | 0.26 | 0.72 |
| detoxification | **Gstm4** | -0.37 | -0.05 | 0.06 | 0.37 |
| detoxification | Gstm5 | -0.34 | 0.12 | 0.26 | 0.70 |
| detoxification | Gstm7 | -0.93 | -0.25 | -0.06 | 0.61 |
| caspases | Casp12 | 0.75 | 0.84 | -0.07 | 0.02 |
| caspases | Casp2 | 0.24 | 0.02 | 0.01 | -0.23 |
| caspases | Casp3 | 1.34 | 0.40 | -0.50 | -1.44 |
| caspases | Casp4 | 0.93 | 0.60 | -0.95 | -1.30 |
| caspases | Casp6 | -0.09 | 0.09 | 0.04 | 0.22 |
| caspases | Casp7 | 0.70 | 0.37 | -0.25 | -0.59 |
| caspases | Casp8 | 1.01 | 0.42 | -0.62 | -1.21 |
| caspases | Casp9 | 0.13 | 0.23 | 0.24 | 0.32 |
| | Acta1 | 3.87 | 2.20 | 0.66 | -1.03 |
| | Myh6 | -0.45 | -0.45 | 0.08 | 0.07 |

$Log_2$

**Supplementary Figure 3. Caspases and xenobiotic gene expression inversely correlates with that of Foxo3 during pressure overload on the heart.** The hearts of mice fed an **A.** IleR or **B.** CR diet before subjecting them to sham or TAC surgeries, were analyzed by RNA-Seq. The heatmap shows the differential RNA-Seq analysis (log2 fold change of the fpkm) of Foxo3, Sod2, Gstm isoforms, and caspases. Numbers that are in black v. grey have a padj of ≤0.05 for the $Log_2$ fold change of TAC/Sham, $TAC^{IleR}/Sham^{IleR}$, $Sham^{IleR}/Sham$, $TAC^{IleR}/TAC$, listed on top of each column.

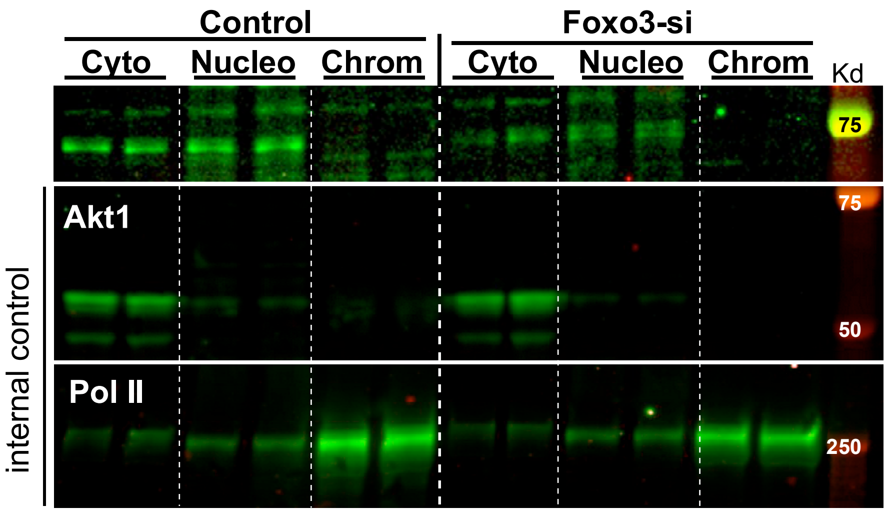

**Supplementary Figure 4**. Knockdown of Foxo3a using adenovirus (Ad5)-delivered shRNA (Foxo3-si). Neonatal cardiac myocytes were infected with 20 moi of a control virus or Ad.Foxo3-si, for 24 h. Cells were harvested, protein fractionated, and analyzed by WB (n=2, each).

A. IleR

| Gene name | TAC/Sham | $TAC^{IleR}/Sham^{IleR}$ | $Sham^{IleR}/Sham$ | $TAC^{IleR}/TAC$ |
|---|---|---|---|---|
| H2-Aa | 0.82 | 0.07 | -1.57 | -2.34 |
| H2-Ab1 | 0.85 | 0.28 | -1.80 | -2.38 |
| **H2-D1** | 0.55 | 0.06 | -0.74 | -1.24 |
| H2-DMb1 | 1.16 | 0.43 | -1.70 | -2.47 |
| H2-Eb1 | 0.83 | 0.17 | -1.74 | -2.42 |
| **H2-K1** | 0.62 | 0.04 | -0.55 | -1.15 |
| H2-M3 | 0.49 | -0.01 | -0.22 | -0.74 |
| H2-Q4 | 0.62 | -0.13 | -0.41 | -1.18 |
| H2-Q5 | 0.78 | -0.24 | -0.75 | -1.79 |
| H2-Q6 | 0.92 | -0.03 | -0.77 | -1.72 |
| H2-Q7 | 1.26 | -0.07 | -0.71 | -2.05 |
| H2-T10 | 1.04 | -0.58 | -0.14 | -1.77 |
| H2-T22 | 0.71 | -0.02 | -0.46 | -1.21 |
| H2-T23 | 0.42 | 0.08 | -0.37 | -0.72 |
| H2-T24 | 0.12 | 0.22 | -1.19 | -1.11 |

B. CR

| Gene name | TAC/Sham | $TAC^{CR}/Sham^{CR}$ | $Sham^{CR}/Sham$ | $TAC^{CR}/TAC$ |
|---|---|---|---|---|
| H2-Aa | 0.87 | 0.52 | -0.01 | -0.36 |
| H2-Ab1 | 0.93 | 0.57 | -0.04 | -0.40 |
| H2-D1 | 0.64 | 0.33 | -0.25 | -0.57 |
| H2-DMb1 | 0.29 | 0.84 | -0.68 | -0.14 |
| H2-Eb1 | 0.85 | 0.39 | -0.03 | -0.50 |
| H2-K1 | 0.66 | 0.23 | -0.04 | -0.48 |
| H2-M3 | 0.49 | 0.13 | -0.18 | -0.54 |
| H2-Q4 | 0.63 | 0.00 | 0.22 | -0.42 |
| H2-Q5 | 0.82 | 0.73 | -0.66 | -0.75 |
| H2-Q6 | 0.76 | 0.10 | -0.18 | -0.85 |
| H2-Q7 | 1.22 | 0.04 | 0.05 | -1.15 |
| H2-T10 | 0.88 | 0.42 | -0.07 | -0.53 |
| H2-T22 | 0.68 | 0.17 | 0.08 | -0.44 |
| H2-T23 | 0.38 | 0.12 | 0.12 | -0.15 |
| H2-T24 | 0.17 | -0.13 | -0.30 | -0.60 |

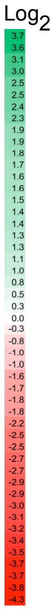

**Supplementary Figure 5. The upregulation of histocompatibility genes after TAC and their suppression by IleR and CR diets.** The hearts of mice fed an **A.** IleR or **B.** CR diet before subjecting them to sham or TAC surgeries, were analyzed by RNA-Seq. The heatmap shows the differential RNA-Seq analysis (log2 fold change of the fpkm) of histocompatibility genes. Numbers that are in black v. grey have a padj of ≤0.05 for the $Log_2$ fold change of TAC/Sham, $TAC^{IleR}/Sham^{IleR}$, $Sham^{IleR}/Sham$, $TAC^{IleR}/TAC$, listed on top of each column.

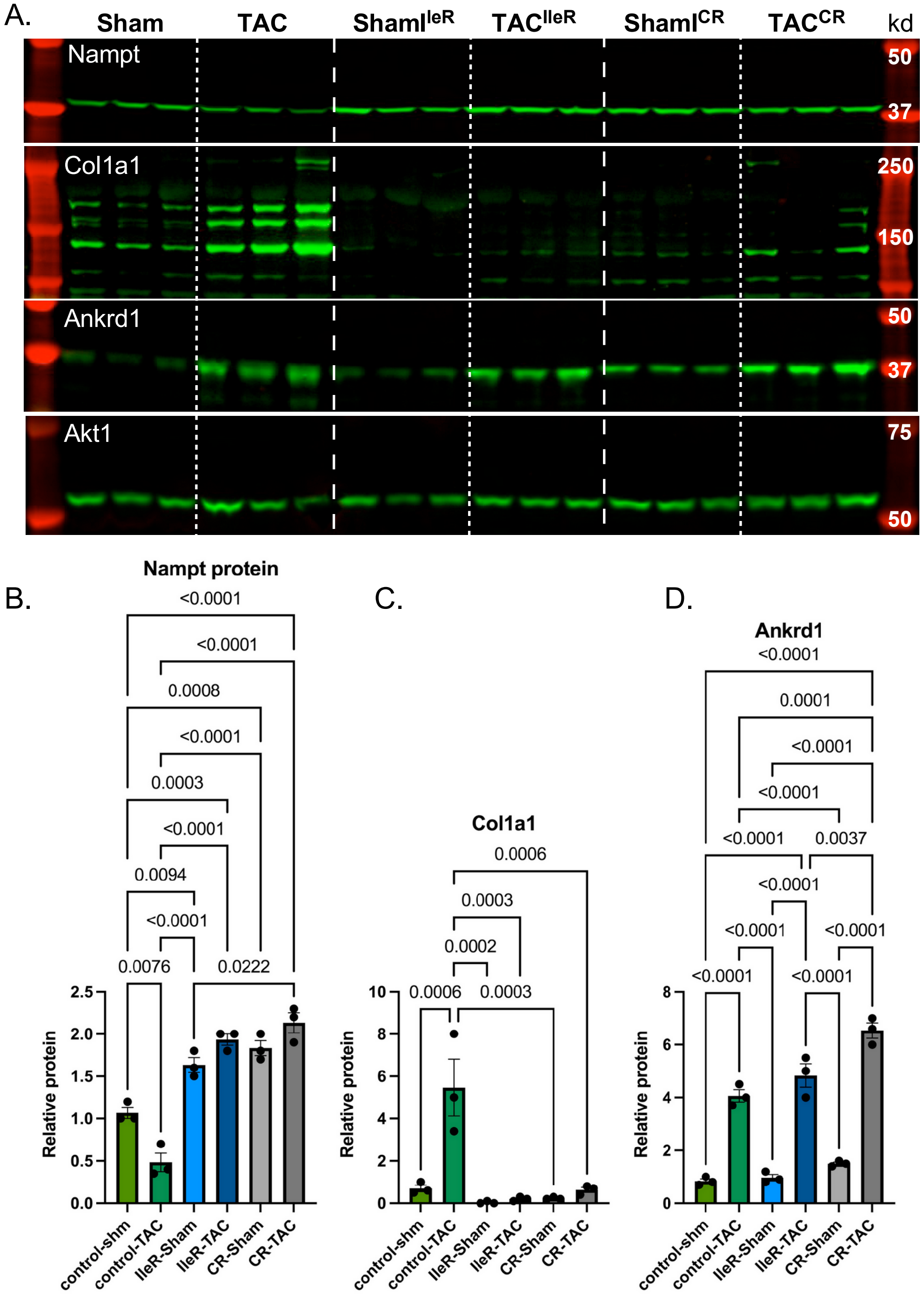

**Supplementary Figure 6. IleR and CR diets inhibit the expression of Col1a1, but not the hypertrophy marker Ankrd1, after 1W of TAC**. Mice were fed a control, IleR, or CR diet 4 days before they were subjected to a sham or TAC surgery and maintained on the same diet for one week. **A-D.** Protein was extracted from the hearts and analyzed by WB (n=3, each). The WB signals were quantitated, normalized to internal control, averaged, graphed, and analyzed by one-way ANOVA (n=3).

A. IleR

| Gene name | TAC/Sham | TAC$^{IleR}$/Sham$^{IleR}$ | Sham$^{IleR}$/Sham | TAC$^{IleR}$/TAC |
|---|---|---|---|---|
| Mylk4 | -1.58 | -0.03 | 0.58 | 2.12 |
| Mylk3 | -0.81 | -0.18 | 0.56 | 1.18 |
| Tcap | -0.74 | 0.07 | 0.78 | 1.58 |
| Tnni3k | -0.69 | 0.00 | -0.02 | 0.65 |
| Tnni3 | -0.64 | -0.06 | 0.32 | 0.89 |
| Tnnt1 | -0.54 | 0.19 | -0.30 | 0.42 |
| Myh6 | -0.43 | 0.04 | 0.15 | 0.60 |
| Myom2 | -0.35 | -0.19 | 1.03 | 1.18 |
| Tnnt2 | -0.31 | -0.10 | 0.83 | 1.03 |
| Myl2 | -0.28 | 0.08 | 0.29 | 0.64 |
| Myl3 | -0.19 | 0.17 | 0.15 | 0.49 |
| Hand2 | -0.17 | -0.22 | 0.60 | 0.54 |
| Ttn | -0.11 | 0.19 | -0.25 | 0.03 |
| Myoz2 | 0.07 | 0.25 | 0.35 | 0.50 |
| Mybpc3 | 0.14 | 0.31 | 0.11 | 0.26 |
| Myom1 | 0.19 | 0.24 | -0.01 | 0.02 |
| Actc1 | 0.23 | 0.43 | -0.34 | -0.16 |
| Tnnc1 | 0.29 | 0.33 | 0.16 | 0.19 |
| Myh7 | 1.17 | -0.08 | 1.09 | -0.17 |
| Acta1 | 3.84 | 1.88 | -0.83 | -2.80 |

B. CR

| Gene name | TAC/Sham | TAC$^{CR}$/Sham$^{CR}$ | Sham$^{CR}$/Sham | TAC$^{CR}$/TAC |
|---|---|---|---|---|
| Mylk4 | -1.60 | -0.76 | 0.14 | 0.97 |
| Mylk3 | -0.82 | -0.37 | 0.71 | 1.15 |
| Tcap | -0.71 | 0.02 | 0.77 | 1.49 |
| Tnni3k | -0.68 | -0.30 | -0.09 | 0.28 |
| Tnni3 | -0.65 | -0.12 | 0.17 | 0.69 |
| Tnnt1 | -0.69 | -0.13 | 0.13 | 0.68 |
| Myh6 | -0.45 | -0.45 | 0.08 | 0.07 |
| Myom2 | -0.35 | -0.23 | 1.47 | 1.58 |
| Tnnt2 | -0.36 | 0.07 | 0.63 | 1.05 |
| Myl2 | -0.30 | -0.01 | 0.47 | 0.75 |
| Myl3 | -0.17 | 0.14 | 0.25 | 0.54 |
| Hand2 | -0.25 | -0.01 | 0.80 | 1.03 |
| Ttn | -0.11 | 0.00 | -0.48 | -0.37 |
| Myoz2 | 0.10 | 0.12 | 0.44 | 0.45 |
| Mybpc3 | 0.16 | 0.24 | 0.21 | 0.28 |
| Myom1 | 0.12 | 0.06 | 0.02 | -0.04 |
| Actc1 | 0.23 | 0.11 | -0.44 | -0.57 |
| Tnnc1 | 0.31 | 0.28 | 0.24 | 0.20 |
| Myh7 | 1.17 | 0.83 | 2.86 | 2.51 |
| Acta1 | 3.87 | 2.20 | 0.66 | -1.03 |

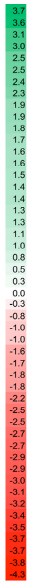

**Supplementary Figure 7. IleR and CR diets increase sarcomeric proteins and the cardiac transcription factor Hand2.** The hearts of mice fed an **A.** IleR or **B.** CR diet before subjecting them to sham or TAC surgeries, were analyzed by RNA-Seq. The heatmap shows the differential RNA-Seq analysis (log2 fold change of the fpkm) of sarcomeric genes. Numbers that are in black v. grey have a padj of ≤0.05 for the Log$_2$ fold change of TAC/Sham, TAC$^{IleR}$/Sham$^{IleR}$, Sham$^{IleR}$/Sham, TAC$^{IleR}$/TAC, listed on top of each column.

### A12450K, A23092901, A23100201 and A23012701-04

#### Rodent Diet w/ 10 kcal% Fat and Same With BCAA Modifications

Formulated by
Research Diets, Inc
Dushyant Kshatriya
10/2/2023

| Product # | A12450K | A23012701 | A23012702 | A23012703 | A23092901 | A23100201 | A23012704 |
|---|---|---|---|---|---|---|---|
| | Control | w/o Leu | w/o Val | w/o Ile | w/ 0.76 g Ile /kg | w/ 1.52 g Ile /kg | w/o Val and Ile |
| **Ingredient** | **gm** | **gm** | **gm** | **gm** | **gm** | **gm** | **gm** |
| Casein | 0 | 0 | 0 | 0 | 0 | 0 | 0 |
| L-Cystine | 4.2 | 4.2 | 4.2 | 4.2 | 4.2 | 4.2 | 4.2 |
| **L-Isoleucine** | **7.6** | **7.6** | **7.6** | **0** | **0.79** | **1.58** | **0** |
| **L-Leucine** | **15.8** | **0.0** | **15.8** | **15.8** | **15.8** | **15.8** | **15.8** |
| L-Lysine | 13.2 | 13.2 | 13.2 | 13.2 | 13.2 | 13.2 | 13.2 |
| L-Methionine | 5.1 | 5.1 | 5.1 | 5.1 | 5.1 | 5.1 | 5.1 |
| L-Phenylalanine | 8.4 | 8.4 | 8.4 | 8.4 | 8.4 | 8.4 | 8.4 |
| L-Threonine | 7.2 | 7.2 | 7.2 | 7.2 | 7.2 | 7.2 | 7.2 |
| L-Tryptophan | 2.1 | 2.1 | 2.1 | 2.1 | 2.1 | 2.1 | 2.1 |
| **L-Valine** | **9.3** | **9.3** | **0.0** | **9.3** | **9.3** | **9.3** | **0.0** |
| L-Histidine | 4.6 | 4.6 | 4.6 | 4.6 | 4.6 | 4.6 | 4.6 |
| L-Alanine | 5.1 | 5.1 | 5.1 | 5.1 | 5.1 | 5.1 | 5.1 |
| L-Arginine | 6.0 | 6.0 | 6.0 | 6.0 | 6.0 | 6.0 | 6.0 |
| L-Aspartic Acid | 12.1 | 12.1 | 12.1 | 12.1 | 12.1 | 12.1 | 12.1 |
| L-Glutamic Acid | 38.2 | 38.2 | 38.2 | 38.2 | 38.2 | 38.2 | 38.2 |
| Glycine | 3.0 | 3.0 | 3.0 | 3.0 | 3.0 | 3.0 | 3.0 |
| L-Proline | 17.8 | 17.8 | 17.8 | 17.8 | 17.8 | 17.8 | 17.8 |
| L-Serine | 10.0 | 10.0 | 10.0 | 10.0 | 10.0 | 10.0 | 10.0 |
| L-Tyrosine | 9.2 | 9.2 | 9.2 | 9.2 | 9.2 | 9.2 | 9.2 |
| Corn Starch | 550 | 565.8 | 559.3 | 557.6 | 556.84 | 556.2 | 566.9 |
| Maltodextrin 10 | 150 | 150 | 150 | 150 | 150 | 150 | 150 |
| Sucrose | 0 | 0 | 0 | 0 | 0 | 0 | 0 |
| Cellulose | 50 | 50 | 50 | 50 | 50 | 50 | 50 |
| Soybean Oil | 25 | 25 | 25 | 25 | 25 | 25 | 25 |
| Lard | 20 | 20 | 20 | 20 | 20 | 20 | 20 |
| Mineral Mix S10026 | 10 | 10 | 10 | 10 | 10 | 10 | 10 |
| DiCalcium Phosphate | 13 | 13 | 13 | 13 | 13 | 13 | 13 |
| Calcium Carbonate | 5.5 | 5.5 | 5.5 | 5.5 | 5.5 | 5.5 | 5.5 |
| Potassium Citrate, 1 H2O | 16.5 | 16.5 | 16.5 | 16.5 | 16.5 | 16.5 | 16.5 |
| Sodium BiCarbonate | 7.5 | 7.5 | 7.5 | 7.5 | 7.5 | 7.5 | 7.5 |
| Vitamin Mix V10001 | 10 | 10 | 10 | 10 | 10 | 10 | 10 |
| Choline Bitartrate | 2 | 2 | 2 | 2 | 2 | 2 | 2 |
| FD&C Yellow Dye #5 | 0 | 0.05 | 0 | 0.025 | 0 | 0 | 0.025 |
| FD&C Red Dye #40 | 0.025 | 0 | 0 | 0.025 | 0.05 | 0 | 0 |
| FD&C Blue Dye #1 | 0.025 | 0 | 0.05 | 0 | 0 | 0 | 0.025 |
| **Total** | **1038.45** | **1038.45** | **1038.45** | **1038.45** | **1038.48** | **1038.58** | **1038.45** |
| **Diet #** | **A12450K** | **A23012701** | **A23012702** | **A23012703** | **A23092901** | **A23100201** | **A23012704** |
| gm | | | | | | | |
| Protein | 178.9 | 163.1 | 169.6 | 171.3 | 172.1 | 172.9 | 162.0 |
| Carbohydrate | 710.0 | 725.8 | 719.3 | 717.6 | 716.8 | 716.2 | 726.9 |
| Fat | 45.0 | 45.0 | 45.0 | 45.0 | 45.0 | 45.0 | 45.0 |
| Fiber | 50.0 | 50.0 | 50.0 | 50.0 | 50.0 | 50.0 | 50.0 |
| gm% | | | | | | | |
| Protein | 17.2 | 15.7 | 16.3 | 16.5 | 16.6 | 16.6 | 15.6 |
| Carbohydrate | 68.4 | 69.9 | 69.3 | 69.1 | 69.0 | 69.0 | 70.0 |
| Fat | 4.3 | 4.3 | 4.3 | 4.3 | 4.3 | 4.3 | 4.3 |
| Fiber | 4.8 | 4.8 | 4.8 | 4.8 | 4.8 | 4.8 | 4.8 |
| kcal | | | | | | | |
| Protein | 716 | 652 | 678 | 685 | 688 | 692 | 648 |
| Carbohydrate | 2840 | 2903 | 2877 | 2870 | 2867 | 2865 | 2908 |
| Fat | 405 | 405 | 405 | 405 | 405 | 405 | 405 |
| Total | 3961 | 3961 | 3961 | 3961 | 3961 | 3961 | 3961 |
| kcal% | | | | | | | |
| Protein | 18 | 16 | 17 | 17 | 17 | 17 | 16 |
| Carbohydrate | 72 | 73 | 73 | 72 | 72 | 72 | 73 |
| Fat | 10 | 10 | 10 | 10 | 10 | 10 | 10 |
| Total | 100 | 100 | 100 | 100 | 100 | 100 | 100 |
| ***Isoleucine (g/kg Diet)*** | ***7.32*** | ***7.32*** | ***7.32*** | ***0.00*** | ***0.76*** | ***1.52*** | ***0.00*** |
| kcal / gm | 3.8 | 3.8 | 3.8 | 3.8 | 3.8 | 3.8 | 3.8 |

Research Diets, Inc.
20 Jules Lane
New Brunswick, NJ 08901 USA


AbdellatifM07

Supplementary Figure 8
